# PIKI-1, a class II phosphatidylinositol 3-kinase, functions in endocytic trafficking

**DOI:** 10.1101/2025.05.22.655458

**Authors:** Gabrielle R. Reimann, Philip T. Edeen, Sylvia Conquest, Barth D. Grant, David S. Fay

## Abstract

Membrane trafficking, including endocytosis and exocytosis, is a complex process that is coordinated by trafficking-associated proteins, cargo molecules, the cytoskeleton, and membrane lipid composition. The NIMA-related kinases NEKL-2 (human NEK8/9) and NEKL-3 (human NEK6/7) are conserved regulators of membrane trafficking in *Caenorhabditis elegans* and are required for successful molting. Through a genetic approach, we isolated reduction-of-function mutations in *piki-1* that suppress *nekl-*associated molting defects. *piki-1* encodes the sole predicted *C. elegans* Class II phosphatidylinositol 3-kinase (PI3Ks), an understudied class of lipid modifiers that contribute to the production of phosphatidylinositol 3-phosphate (PI(3)P) and phosphatidylinositol 3,4-bisphosphate (PI(3,4)P_2_). Using a set of genetically encoded lipid sensors, we found that PIKI-1 was responsible for the production of PI(3,4)P_2_ in the *C. elegans* epidermis but played only a minor role in the control of PI(3)P levels. Consistent with this, both PI(3,4)P_2_ and PIKI-1 colocalized to early endosomes, and reduction of PIKI-1 function strongly affected early endosomal morphology and protein composition. Additionally, reduced PIKI-1 function led to excess tubulation of endosomal compartments associated with recycling or the degradation of cellular debris. In contrast to previous studies using mammalian cell culture, PIKI-1 was largely dispensable for clathrin-mediated endocytosis in the context of the worm epidermis, which is a polarized epithelium. Notably, reduction of PIKI-1 function strongly mitigated defects in early endosomes associated with the depletion of NEKL-2. We propose that reduction of PIKI-1 function may suppress *nekl* molting defects by partially restoring endocytic trafficking within specific compartments, including the early endosome. We also show that inhibition of the PI(3,4)P_2_-binding protein HIPR-1 (HIP1/HIPR1) suppresses *nekl* molting defects, suggesting that reduced PI(3,4)P_2_ levels alter endosomal protein recruitment in a manner that antagonizes NEKL-2 function.

**Author summary:** The uptake of materials from outside the cell and their subsequent delivery to specific intracellular locations are essential for cell function and survival. Two of the mechanisms that control this complex intracellular pathway involve the modification of proteins and of lipids, processes that are highly conserved across species. In this study, we used the model organism *Caenorhabditis elegans*, which is highly amenable to cell biological and genetic approaches, to establish a novel connection between these two regulatory mechanisms and demonstrate the importance of lipid modifications in maintaining the normal functioning of intracellular transport. Our results also provide insights into the fundamental cellular functions of proteins associated with human disease including cancer and metabolic disease.

## Introduction

Embedded in the cytosolic leaflet of cellular membranes are small phospholipids known as phosphoinositides [1–4]. Phosphoinositides have a glycerol backbone esterified to two fatty acid chains and a phosphate and a polar head group, termed *myo*-inositol, that extend into the cytoplasm (Fig 1A) [1–4]. Modification of the *myo*-inositol head group at hydroxyl positions 3’, 4’, and 5’ generates seven different phosphoinositide species; conversion between species is controlled in a spatiotemporal manner by different classes of lipid phosphatases and kinases (Fig 1B) [1–5]. Phosphoinositides make up only ∼10% of the total membrane phospholipid pool in the cell and are comparatively short-lived [1]. Nevertheless, the concentration of specific phosphoinositides displayed on the cytoplasmic leaflet of an organellar membrane controls the recruitment and activity of trafficking and signaling regulators, thereby impacting key cellular processes [4–7].

**Fig 1.**
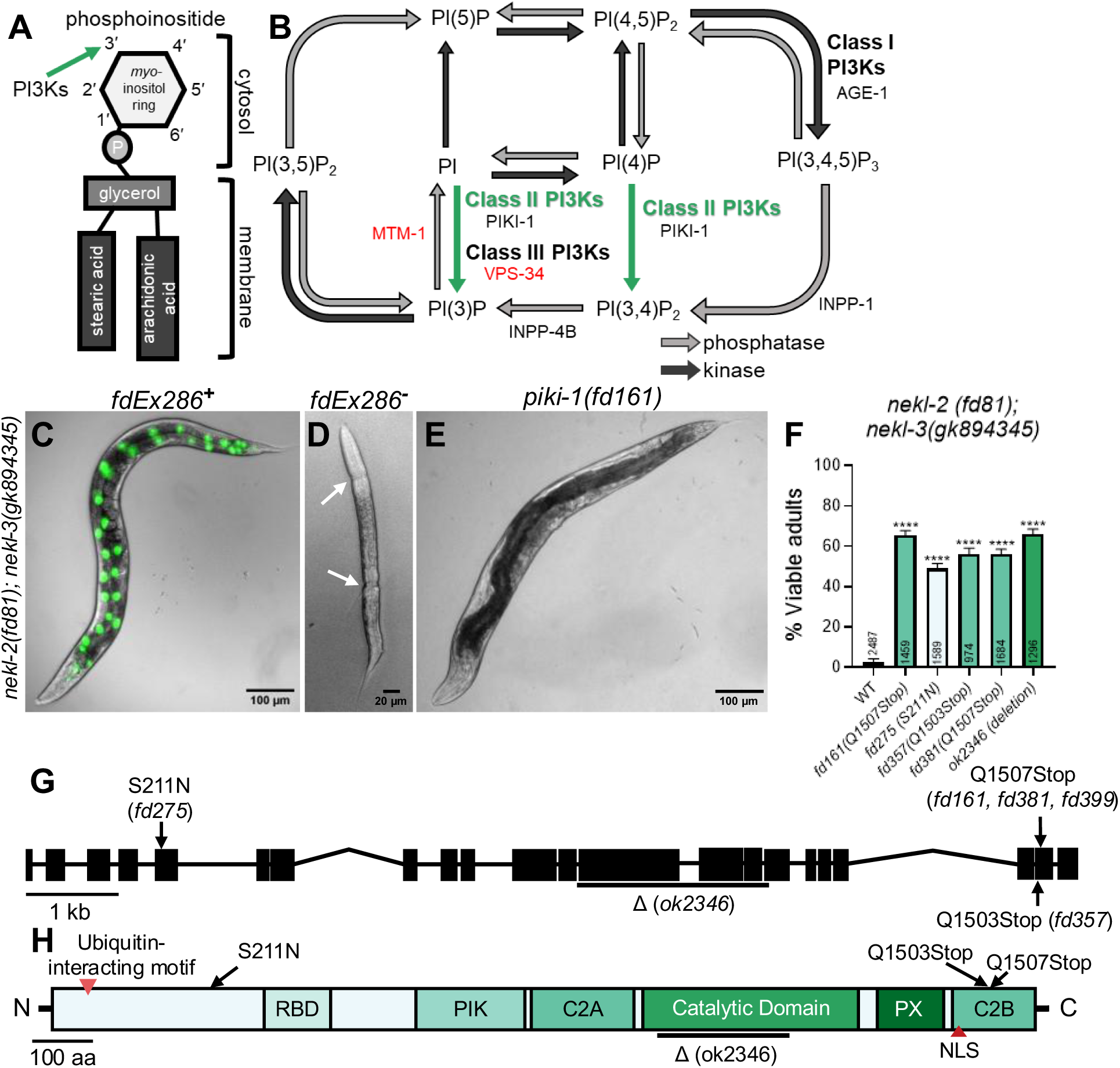
*nekl*-associated molting defects are suppressed by loss of function of *piki-1*. (A) Graphical representation of the basic structure of a phosphoinositide The 3’ position in the inositol ring that is phosphorylated by phosphatidylinositol 3-kinases (PI3Ks) is indicated by the green arrow. (B) Schematic diagram of the phosphoinositide biosynthesis pathway. Dark gray arrows represent lipid kinase activity, and light gray arrows indicate lipid phosphatase activity. The reactions catalyzed by the different classes of PI3Ks are indicated. The green arrows indicate reactions catalyzed by class II PI3Ks. *C. elegans* genes coding for kinases and phosphatases are listed by the reaction they catalyze. Essential genes are in red text. (C) Merged DIC and fluorescence image of a representative *nekl-2(fd81); nekl-3(gk894345)* worm that has an extrachromosomal rescuing array (*fdEx286*). *fdEx286* contains wild-type *nekl-3* and a broadly expressed reporter (SUR-5::GFP), seen in green in the image. (D) DIC image of an arrested *nekl-2(fd81); nekl-3(gk894345)* larva that did not inherit *fdEx286*. White arrows indicate cuticular boundaries, where the old cuticle has been shed from the head and tail but not from the midbody. (E) Representative DIC image of a suppressed *nekl-2(fd81); nekl-3(gk8942345); piki-1(fd161)* mutant, which does not require *fdEx286* to propagate. (F) The percentage of viable adults for the indicated genotypes. The corresponding amino acid changes are indicated in parentheses next to the gene. The number of animals analyzed per genotype is indicated in the graph. Data are shown as the mean and 95% confidence interval (CI). Statistical significance was determined using unpaired t-tests; ****p ≤ 0.0001 relative to control. Raw data available in S1 File. (G) Schematic diagram of the *piki-1* gene. Black rectangles denote exons and thin lines denote introns. Locations of missense alleles are indicated by solid arrows. The large deletion (Δ) *ok2346* is indicated by a line beneath the affected exons and introns. (H) Schematic diagram of the PIKI-1 protein. Red arrowheads indicate known motifs. Black arrows indicate the mutated amino acids. A black line indicates the region affected by the ok2346 deletion. C2A, first C2 domain; C2B, second C2 domain; [NLS, nuclear localization signal;] PIK, PI3K class II accessory domain; PX, phox homology domain; RBD, Ras**-**binding domain.

The interconnected nature of the phosphoinositide pathway and the ability of phosphoinositides to undergo reversible modifications by lipid modifiers allow for precise control of the phosphoinositides on cellular membranes [5–7]. One family of lipid modifiers phosphatidylinositol 3-kinases (PI3Ks), which specifically phosphorylate the hydroxyl group at position 3’ on the inositol ring to produce three different phosphoinositide species: phosphatidylinositol (3,4,5)-trisphosphate [PI(3,4,5)P_3_], phosphatidylinositol 3,4-biphosphate [PI(3,4)P_2_], and phosphatidylinositol 3-phosphate [PI(3)P] (Fig 1B) [8–11]. PI3Ks are subdivided into three classes, based on the phosphoinositide species they produce, their protein domains, and their interactions with different regulatory subunits [10,11]. Class I PI3Ks produce PI(3,4,5)P_3_ and are involved in multiple pathways that regulate cell growth, proliferation, metabolism, and autophagy [12,13]. Class III PI3Ks produce PI(3)P and primarily regulate membrane trafficking, most notably endosome-to-lysosome maturation and autophagy [12,13]. Class II PI3Ks are not as well characterized as Class I or III PI3Ks, due in part to the overlap in their contributions to phosphoinositide production [10–13]. Class II PI3Ks can produce both PI(3)P and PI(3,4)P_2_ [6,14–16]. PI(3)P can also be produced by Class III PI3Ks [6,17,18], whereas PI(3,4)P_2_ can be generated by the dephosphorylation of the Class I PI3K product, PI(3,4,5)P_3_ (Fig 1B) [19–24]. Class II PI3Ks also differ from Class I and III PI3Ks in that they do not form complexes with additional regulatory units [14,25–30].

The role of PI3Ks and their products is critical for multiple steps in membrane trafficking [1,3,31]. For example, uptake of cargo by clathrin-mediated endocytosis is dependent on phosphatidylinositol 4,5-bisphosphate [PI(4,5)P_2_], which is required for initiating formation of the clathrin lattice on budding vesicles [32–36]. During maturation of the clathrin-coated pit, PI3Ks and lipid phosphatases are subsequently recruited to modify the phosphoinositide population, which is necessary for the recruitment of proteins that promote vesicle scission and the internalization of cargo [37]. Production of another phosphoinositide species, PI(3,4)P_2_, which is derived from modification of PI(4) by class II PI3Ks via PI(4,5)P_2_, is implicated in the recruitment of proteins that promote scission of the vesicle [7,15,21,38,39].

After internalization, nascent vesicles fuse with the early endosome (also referred to as the sorting endosome), where cargo is sorted for delivery to specific intracellular locations [40,41]. Within the early endosome, the phosphoinositide population consists primarily of PI(3)P, produced predominantly by the class III PI3K, with some contribution from class II PI3Ks [42,43]. Here, PI(3)P recruits early endosome effectors necessary for vesicular docking, fusion, and the subsequent sorting of cargo [6,42–45]. During maturation of the early endosome, subdomains develop, a process that is driven by cargo, membrane-associated proteins, and underlying changes in the composition of the phosphoinositides [43]. Phosphorylation by the PI(3)P 5-kinase produces phosphatidylinositol 3,5-bisphosphate [PI(3,5)P_2_], which regulates the sorting of proteins destined for late endosomes and degradation in the lysosome [43,46,47]. The production of PI(3,4)P_2_ on subdomains of the early endosome has been proposed to play a role in recruiting and activating Rab11 and directing the transport of cargo to recycling pathways [48,49]. The route for the production of PI(3,4)P_2_ on the surface of membranes is unclear, but there is emerging evidence that PI(3,4)P_2_ is synthesized directly by class II PI3Ks and is not derived solely from the dephosphorylation of PI(3,4,5)P_3_ [19–22,38,50–52].

One of the challenges of studying PI3Ks and the roles of their lipid products relates to the transient nature of phosphoinositide species [53–55]. Additionally, the pathways regulated by PI3Ks exhibit a level of interdependency that complicates the interpretation of experiments involving enzymatic inhibition by genetic manipulation or the use of selective inhibitors [13]. Moreover, highly selective inhibitors are not available for class II PI3Ks [13,27,56]. Lastly, the ability to visualize phosphoinositides inside cells using antibodies against phosphoinositides has not been widely used because the fixation required precludes live imaging, which is invaluable for understanding the localization, dynamics, and functions of these small lipids [53,57]. This last challenge has been addressed by the development of genetically encoded lipid biosensors, which tether fluorescent proteins to lipid-binding domains that can recognize individual phosphoinositide species [58–60]. Nevertheless, the use of lipid biosensors is complicated by their cross-specificity with different lipids, by difficulties with matching sensor levels to the abundance of the phosphoinositide species, and by potential physiological effects that the sensor may cause by competing with native phosphoinositide-binding proteins for lipid-binding sites [53,58–60]. Despite these complications, lipid biosensors are currently the best tool available for the visualization of intracellular phosphoinositide pools and for understanding their roles in membrane trafficking.

Here, we use the small nematode, *Caenorhabditis elegans*, as a model for studying endocytic trafficking within an intact organism [61]. Endocytic trafficking is closely coupled to the *C. elegans* molting cycle, as the inhibition of membrane trafficking proteins can lead to strong defects in the molting process [62]. Previously, we have shown that two conserved NIMA-related Ser/Thr protein kinases, NEKL-2 (NEK8/9 in mammals) and NEKL-3 (NEK6/7), and their binding partners, the conserved ankyrin repeat proteins MLT-2, MLT-3, and MLT-4 (mammalian ANKS6, ANKS3, and INVS, respectively), are essential for proper molting via the regulation of endocytic trafficking [62–67]. We have previously described a genetic approach for identifying suppressors of *nekl-*associated molting defects [68]. These screens led to the identification of several conserved regulators of membrane trafficking, including TAT-1 (mammalian ATP8A1/2), a phosphatidylserine flippase, which controls lipid asymmetry on the surface of recycling endosomes [69]. Here we report the identification of mutations affecting PIKI-1, a *C. elegans* ortholog of mammalian class II PI3Ks, which comprise three family members (PI3KC2A, PI3KC2B, PI3KC2G) [8,10,12,14]. Importantly, PIKI-1 is the sole predicted class II PI3K in *C. elegans*, which allowed us to study class II PI3K functions in the absence of genetic redundancy. We found that reduction of PIKI-1 activity led to defects in several endocytic compartments, most notably the early endosome. To understand the cellular functions of PIKI-1, we generated several phosphoinositide biosensors to visualize phosphoinositide pools in the epidermis of adult worms. Our study indicated that whereas PIKI-1 is a minor contributor to PI(3)P pools in the epidermis, it is a major contributor to PI(3,4)P_2_ levels, supporting the model that PI(3,4)P_2_ is predominantly synthesized by class II PI3Ks and not through the degradation of class I PI3K– derived products.

## Results

### *nekl* reduction-of-function molting defects are suppressed by reduction-of-function mutations in *piki-1*

We previously described a forward genetic screen coupled with a whole-genome sequencing pipeline to identify genetic suppressors of larval lethality caused by *nekl* reduction-of-function mutations [68]. Briefly, this approach uses a synthetic lethal interaction that occurs when two aphenotypic reduction-of-function alleles of *nekl-2(fd81)* and *nekl-3(gk894345)* are combined in the same animal. Consistent with our earlier studies [63], *nekl-2(fd81)*; *nekl-3(gk894345)* (hereafter *nekl-2; nekl-3*) worms showed a highly penetrant developmental defect, with ∼98% of progeny arresting at the L2/L3 boundary and only ∼2% becoming viable adults (Fig. 1D, 1F). *nekl-2; nekl-3* worms were propagated in the presence of a rescuing array (*fdEx286*) containing wild-type copies of *nekl-3* and a *sur-5::gfp* reporter to facilitate the visualization of animals containing the array (Fig 1C). Worms that did not inherit the array were molting defective and exhibited a corset phenotype in which the old cuticle was shed from the anterior and posterior of the worm but not from the midbody (Fig 1D). After mutagenesis, worms containing suppressor mutations were identified by their ability to propagate in the absence of the rescuing array (Fig 1E).

From our suppressor screen, we identified three independent alleles that affect the *piki-*1 locus (*fd161*, *fd275*, and *fd357*), such that ∼50–65% of these *nekl-2; nekl-3 piki-1* worms reached adulthood (Fig 1F). Two alleles, *fd161* and *fd357*, cause a C-to-T transition in the second-to-last exon of *piki-1*, resulting in an early stop codon at amino acids 1507 and 1503, respectively (Fig 1G, 1H). The third allele, *fd275*, is a G-to-A transition in the fifth exon of *piki-1*, leading to the substitution of asparagine for serine at amino acid 211 (Fig 1G, 1H). CRISPR phenocopy of *fd161* (*fd381*) led to levels of suppression that were similar to those observed for *fd161* (Fig 1F), demonstrating that *piki-1* is the causal locus in the *nekl-2; nekl-3 piki-1(fd161)* strain. Additionally, *fd161* failed to complement *fd275*, consistent with *piki-1* being the causal locus in both strains. Moreover, a consortium-generated 1597-bp deletion mutation in *piki-1* (*ok2346*) led to 66% of *nekl-2; nekl-3 piki-1(ok2346)* worms reaching adulthood (Fig 1F–1H). We note, however, that we failed to observe robust suppression of *nekl-2; nekl-3* defects by *piki-1(RNAi)*, which may be due to insufficient knockdown of *piki-1* using this method (4.8%, n=2114; S1 Fig)

PIKI-1 belongs to a family of lipid kinases that specifically phosphorylate the 3’ hydroxyl position on the inositol ring of membrane phosphoinositides (Fig 1A). More specifically, PIKI-1 is the sole member of the class II PI3Ks in *C. elegans*, which in other species have been reported to convert PI to PI(3)P and PI(4)P to PI(3,4)P_2_ (Fig. 1B). Given the proposed role of PIKI-1 in PI modifications, we tested several non-essential PI modifiers for genetic interactions with the *nekls* (Fig 1B). PI(3,4)P_2_ can be derived from PI(3,4,5)P_3_ after removal of the 5’ phosphate (Fig 1B) (refs). We found that a reduction-of-function mutation in *age-1* (*hx546*), the sole class I PI3K in *C. elegans*, failed to suppress *nekl-2; nekl-3* mutants (S1 Fig). Additionally, RNAi-mediated inhibition of *age-1*, *inpp-1* (a 5′ phosphatase targeting PI(3,4,5)P₃), or *inpp-4b* (a 4′ phosphatase targeting PI(3,4)P_2_) failed to alter the molting phenotypes of *nekl* mutants (S1 Fig). Nevertheless, RNAi of *inpp-1* or *inpp4b* led to an ∼70% reduction in brood size in the *nekl-2; nekl-3* background (S1 file), indicating that reduction of INPP-1 or INPP-4b activity may impact fertility. Collectively, these results suggest that the suppression of *nekl* defects by PI-pathway modifiers may be specific to *piki-1*. We note, however, that our inability to test essential PI modifiers and the caveat of partial knockdown by RNAi limit our conclusions.

### PIKI-1 localizes to clathrin-coated pits and early endosomes

Previously, we reported colocalization of NEKL-2 and NEKL-3 with several membrane trafficking compartments, consistent with roles for the NEKLs in endocytic trafficking [66]. More specifically, NEKL-2 localizes primarily to early endosomes, whereas NEKL-3 resides predominantly at late endosomes [66]. Mammalian class II PI3Ks localize both to early endosomal compartments and to clathrin-coated structures, as well as to recycling endosomes [14,70,71]. In *C. elegans*, PIKI-1 localizes to nascent phagosomes in embryos and in the germline [72–76].

To characterize the endogenous expression and localization of PIKI-1 in the epidermis, we initially examined CRISPR-generated fusions of either GFP or mScarlet to the C terminus of PIKI-1. However, endogenous PIKI-1::GFP proved too dim for reliable imaging and PIKI-1::mScarlet showed non-specific localization to lysosomes (S2 Fig), which is likely due to the cleavage of the fluorophore and its retention within lysosomes, as has been reported [77]. We therefore integrated a single-copy PIKI-1::mNeonGreen transgene driven by a hyp7-specific promoter (P_hyp7_; *semo-1*) using miniMos methods [78]. We note that for technical reasons we were unable to obtain a marker for PIKI-1 fused to an acid-sensitive red fluorescent protein. P_hyp7_::PIKI-1::mNeonGreen was detected in the adult epidermis at apical puncta as well as more diffusely throughout the epidermis (Fig 2B, 2B’). Based on the distribution and size of P_hyp7_::PIKI-1::mNeonGreen puncta, we hypothesized that PIKI-1 may localize to clathrin-coated structures and/or apical endosomes, consistent with some prior studies on mammalian class II PI3Ks.

**Fig 2.**
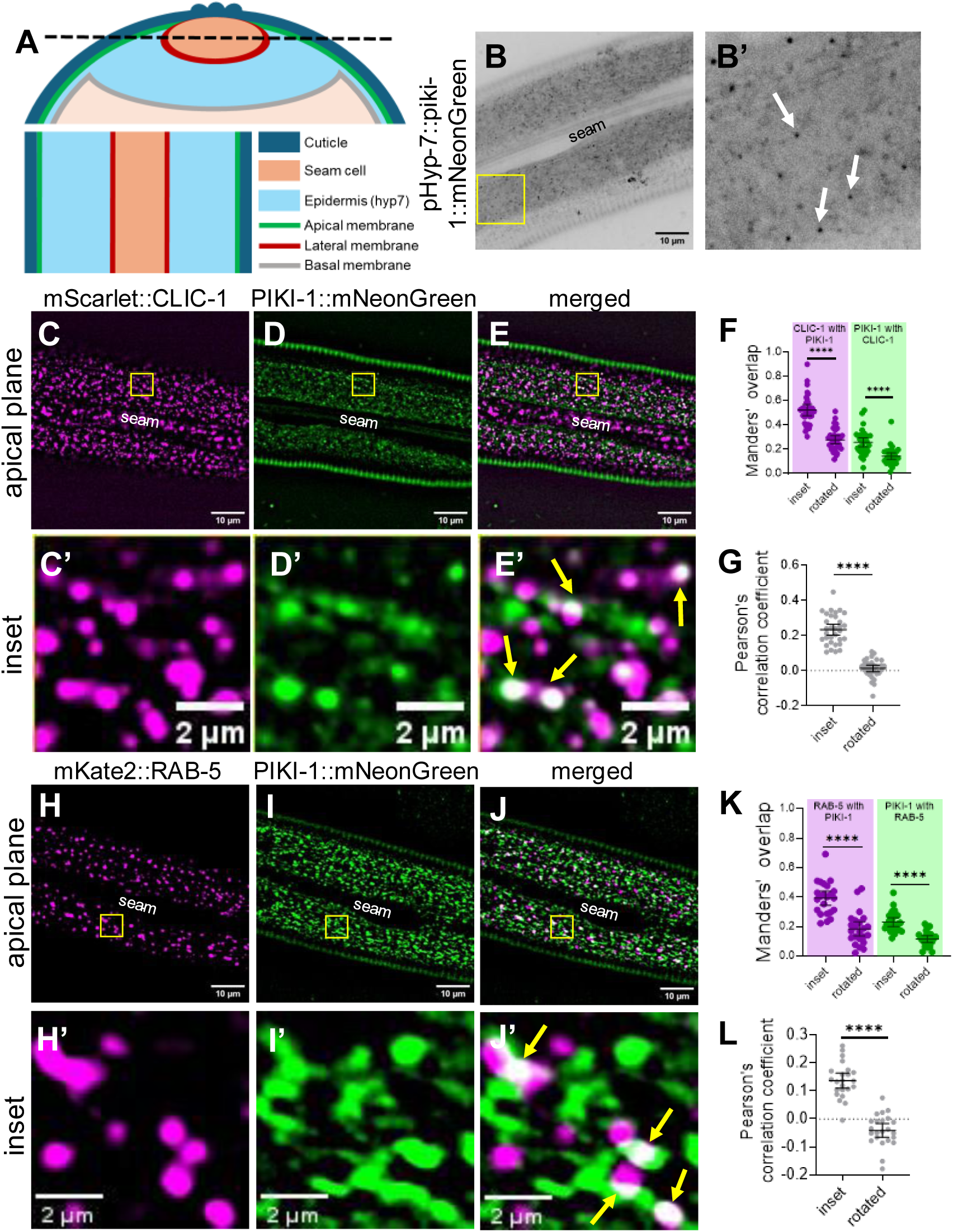
PIKI-1 colocalizes with clathrin-coated pits and the early endosome. (A) Schematic diagram of the adult *C. elegans* as a transverse cross-section (top) and a longitudinal cross-section (bottom), which corresponds to the apical plane. The black dashed line on the transverse cross-section denotes the apical plane visualized in the longitudinal cross-section for a single z-slice. On the bottom right is a key indicating the colors corresponding to different tissues and membranes. (B, B’) Representative image of a Day-1 adult expressing P_hyp7_::PIKI-1::mNeonGreen. White arrows indicate apical puncta. (C-E’) Colocalization assays were carried out in worms expressing (C, C’) CLIC-1::mScarlet and (D, D’) P_hyp7_::PIKI-1::mNeonGreen (n=32); (E, E’) merged images. (H–J’) Trans-heterozygous worms were used for colocalization between (H,H’) P*_dyp-7_*::mKate2::RAB-5 and (I,I’) P_hyp7_::PIKI-1::mNeonGreen (n=23); (J,J’) merged images. The seam cell is labeled in lower-magnification images. Yellow squares (C– D, H–J) indicate enlarged insets (C’–E’, H’–J’). In merged insets (E’ and J’), yellow arrows indicate examples of colocalization. (F, G, K, L) Colocalization was quantified using Mander’s overlap (F, K) and Pearson’s correlation coefficient (G, L). Dot plots show the mean and 95% CI. Statistical significance between rotated and inset values was determined by unpaired *t-*tests; ****p ≤ 0.0001. Raw data available in S1 File.

As anticipated, we observed colocalization between a marker for the clathrin light chain, mScarlet::CLIC-1, and PIKI-1::mNeonGreen. However, whereas ∼50% of the mScarlet::CLIC-1 signal overlapped with the PIKI-1::mNeonGreen signal (Manders’ overlap = ∼0.5), only ∼25% of PIKI-1::mNeonGreen overlapped with mScarlet::CLIC-1 (Manders’ overlap = ∼0.25) (Fig 2C–2F), suggesting that PIKI-1 may localize to additional membrane compartments. Significant positive but partial overlap between the markers was also supported by the Pearson’s correlation coefficient (PCC = ∼0.23) (Fig 2G). As an additional control, we rotated one of the two channels and reanalyzed the Manders’ and Pearson’s values and observed a dramatic reduction in both measurements, indicating that the overlap was non-random (Fig 2F, 2G).

We next examined colocalization between PIKI-1::mNeonGreen and a marker for early endosomes, P*_dpy-7_*::mKate2::RAB-5. We found that 40% of the mKate2::RAB-5 signal overlapped with PIKI-1::mNeonGreen (Manders’ overlap = ∼0.4; Fig 2H–2K), whereas 23% of the PIKI-1::mNeonGreen signal overlapped with mKate2::RAB-5 (Manders’ overlap = ∼0.23; Fig 2H–2K). Additionally, the Pearson’s correlation coefficient (PCC = ∼0.14) is consistent with overlap between mKate2::RAB-5 and PIKI-1::mNeonGreen (Fig 2L). These results indicate that PIKI-1 is associated with both clathrin-coated pits and endosomal trafficking compartments, suggesting that the role of the class II PI3 kinases in endomembrane trafficking is conserved between *C. elegans* and mammals.

### Loss of PIKI-1 affects endosomal compartments

Mammalian class II PI3Ks are implicated in the scission of nascent clathrin-coated vesicles and the subsequent uncoating and maturation of vesicles enroute to the early endosome [70]. To determine whether PIKI-1 regulates early steps in clathrin-mediated endocytosis, we examined markers for clathrin light and heavy chains in wild type and *piki-1* mutants. Despite localization of PIKI-1 to clathrin-coated pits, *piki-1(Q1507Stop)* mutants showed a wild-type-like pattern of localization for both clathrin heavy (GFP::CHC-1) and light (mScarlet::CLIC-1) chains, suggesting that PIKI-1 does not play a major role in early steps of clathrin-mediated endocytosis in the *C. elegans* epidermis (S3 Fig). Consistent with this, LRP-1, an apically expressed low-density lipoprotein–like receptor that is trafficked through clathrin-coated pits (Fig 3A) [79], showed only slightly altered localization at the apical surface in *piki-1(Q1507Stop)* worms as compared with wild type (S3 Fig). Collectively, our data indicate that PIKI-1 may be largely dispensable during clathrin-mediated endocytosis.

**Fig 3.**
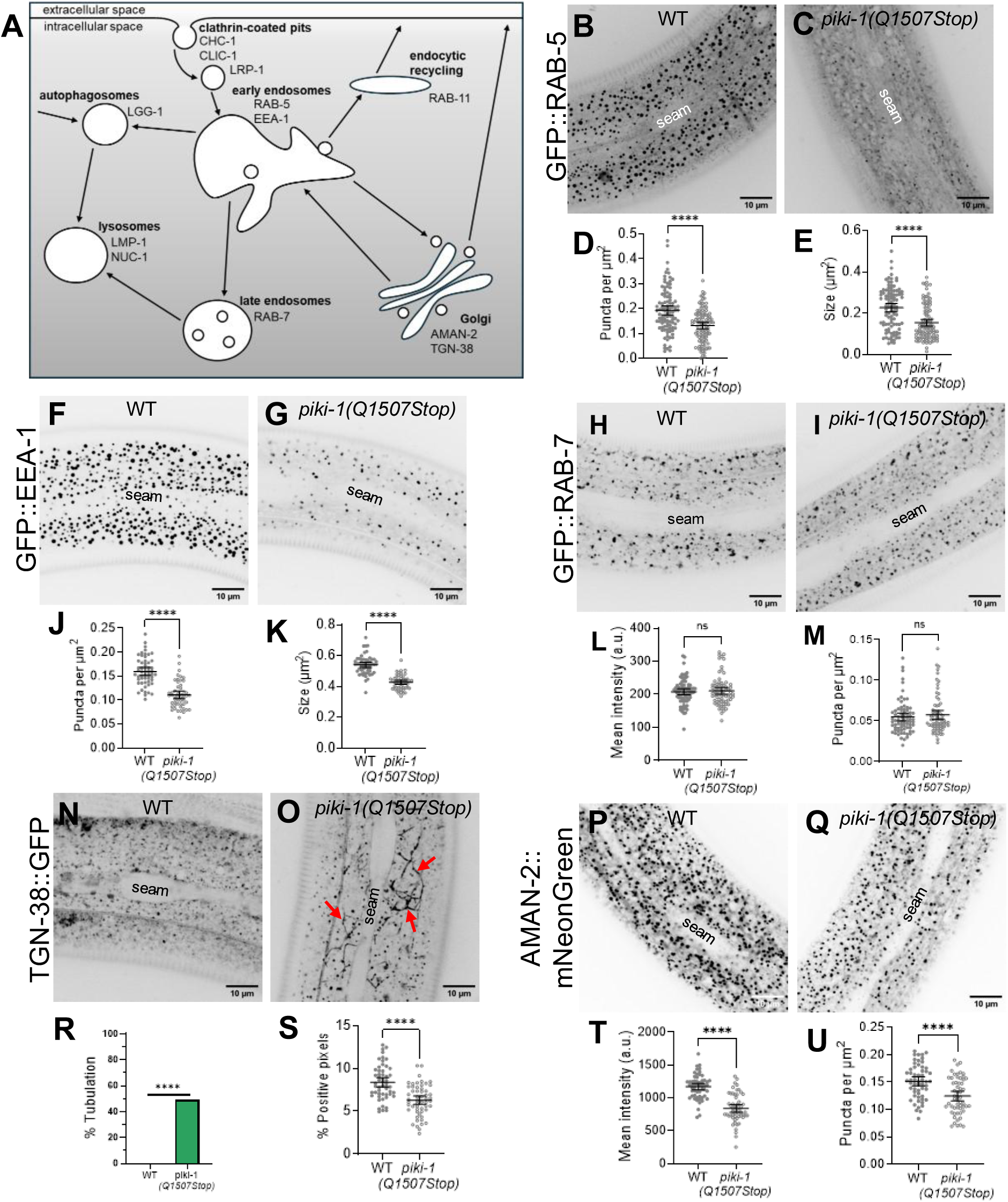
Reduction of PIKI-1 function affects different endosomal compartments. (A) Schematic of endocytosis and associated endosomal compartments. Compartments are labeled in bold. Proteins and cargos affiliated with the compartments are below the bolded compartment labels. (B,C,F,G,H,I,N,O,P,Q) Representative confocal images of day-1 adults that were used to assess the markers (B,C) P*_rab-5_*::GFP::RAB-5 (n = 106), (F, G) P_hyp7_::GFP::EEA-1 (n = 54), (H,I) P_hyp7_::RAB-7 (n = 74), (N, O) P_hyp7_::TGN-38::GFP (n = 55), and (P, Q) P_hyp7_::AMAN-2::mNeonGreen (n = 56) in *piki-1(Q1507Stop)* mutants relative to wild-type worms. The seam cell is labeled in all images. (D,E,J,K) The (D,J) number of puncta per unit area and the (E,K) size of puncta were plotted for worms expressing (D,E) P*_rab-5_*::GFP::RAB-5 and (J,K) P_hyp7_::GFP::EEA-1. (L,M) The (L) mean intensity (arbitrary units, a.u.) and the (M) number of puncta per unit area were plotted for worms expressing P_hyp7_::RAB-7. (R) The percent positive pixels (above threshold) was plotted for worms expressing P_hyp7_::TGN-38::GFP. (S) Worms expressing P_hyp7_::TGN-38::GFP were scored for the presence of tubulations within the epidermis and are graphed to show the percentage of worms that have no tubulation and that have tubulations. (O) Red arrows indicate tubulations. (T, U) The (T) mean intensity and the (U) number of puncta per unit area were plotted for worms expressing P_hyp7_::AMAN-2::mNeonGreen. All dot plots show the mean and 95% CI. Statistical significance was determined by unpaired *t-*tests; ****p ≤ 0.0001; ns, not significant. (R) Statistical significance was determined by Fisher’s exact test; ****p ≤ 0.0001. Raw data are available in S1 File.

After internalization at the plasma membrane, cargoes are next sorted at the early endosome for subsequent routing to either recycling or degradative pathways [40]. To assess the role of PIKI-1 on early endosomes, we examined two early endosomal reporters, GFP::RAB-5 and GFP::EEA-1, in wild type and *piki-1* mutants. RAB-5 is a small GTPase that recruits proteins required for endocytic sorting, whereas EEA-1 is a conserved effector of RAB-5 and promotes vesicle docking and fusion (Fig 3A) [44,45]. Notably, we observed a marked (∼1.5-fold) reduction in the number of RAB-5– and EEA-1–positive endosomes in *piki-1(Q1507Stop)* mutants as compared with wild type (Fig 3B–3D, Fig 3F, 3G, 3J). Moreover, the size of RAB-5– and EEA-1– positive vesicles was reduced by ∼1.5- and ∼1.3-fold, respectively (Fig 3E, 3K). These results indicate that reduction of PIKI-1 function leads to a decrease in the size of the early endosome compartment in the epidermis. In contrast, reduction of PIKI-1 function had no detectable effect on the mean intensity or morphology of late endosomes as marked by GFP::RAB-7 (Fig 3H, 3I, 3L, 3M). These findings are consistent with PIKI-1 playing a role in the function of early endosomes in *C. elegans*.

Other studies have implicated mammalian Class II PI3Ks in regulating cargo recycling pathways [49,71,80]. To evaluate the role of PIKI-1 in endocytic recycling, we first examined the localization pattern of GFP::RAB-11, which marks one of the major recycling compartments. We observed a very slight decrease (∼1.1-fold) in mean intensity of GFP::RAB-11 in *piki-1(Q1507Stop)* mutants (p = 0.0012), as well as a modest (though not statistically significant) decrease (∼1.3-fold; p = 0.15) in the size of the GFP::RAB-11 puncta (S3 Fig). These results suggest that PIKI-1 may have only a minor or indirect role in the function of the GFP::RAB-11 recycling compartment.

We next examined localization of the cargo protein TGN-38, which travels from the Golgi to the plasma membrane during exocytosis and is recycled back to the Golgi via early endosomes, potentially bypassing the RAB-11 compartment [81,82]. Although no changes in mean intensity of TGN-38::GFP were observed between wild type and *piki-1(Q1057Stop)* mutants (S3 Fig), we noticed a striking tubulation phenotype, as marked by TGN-38::GFP, in ∼50% of *piki-1(Q1507Stop)* worms (Fig 3N, 3O, 3R). Consistent with the altered distribution of TGN-38::GFP, we detected a decrease in the percentage of TGN-38::GFP–positive pixels (above threshold) in *piki-1(Q1057Stop)* animals relative to wild type (Fig 3N, 3O, 3S). These findings suggest that PIKI-1 plays a role in endocytic recycling, albeit independently of the RAB-11 recycling pathway.

Given our findings with TGN-38, we assessed the impact of PIKI-1 reduction on the Golgi by using the AMAN-2::mNeonGreen marker [83]. Notably, we observed an ∼1.4-fold reduction in the mean intensity of AMAN-2::GFP in *piki-1(Q1507Stop)* mutants (Fig 3P, 3Q, 3T). Additionally, we detected a modest decrease in the number of AMAN-2 puncta (∼1.2-fold) in *piki-1* mutants (Fig 3P, 3Q, 3U). Taken together, our results point to a role for PIKI-1 at the early endosome and suggest that the sorting or transport of cargo from the early endosome to the Golgi is disrupted in the absence of PIKI-1.

As PIKI-1 promotes autophagy in the developing embryo and in the adult germline [74–76, 84], we next investigated the role of PIKI-1 in autophagy in the adult epidermis. We examined the expression of an autophagosome marker, mNeonGreen::LGG-1 (Fig 3A) [85,86], in wild type and *piki-1* mutants. We observed a modest reduction (∼1.3-fold) in the mean intensity of mNeonGreen::LGG-1 in *piki-1(Q1507Stop)* mutants (S3 Fig). In addition, we observed an increase in the frequency of elongated or tubular structures marked by mNeonGreen::LGG-1 in *piki-1(Q1507Stop)* mutants (38.2%) as compared with wild type (10%) (S3 Fig). This phenotype is similar to what we observed with the TGN-38 marker (Fig 3N, 3O, 3R), although the effect on LGG-1 was less robust. These results suggest that, similar to findings in the embryo and adult germline, PIKI-1 also plays a role in autophagy in the adult epidermis. Taken together, our data support a direct role for PIKI-1 in endocytic trafficking, primarily at the level of the early endosome, with additional effects observed on endocytic recycling and early steps in autophagy.

### Loss of PIKI-1 suppresses defects in early endosomes associated with NEKL-2 depletion

Given the strong effects observed for early endosomes, we were curious whether the role of PIKI-1 at early endosomes could account for the genetic suppression of *nekl-2; nekl-3* mutants. In addition, we had previously observed localization of NEKL-2 to early endosomes and had observed that depletion of NEKL-2 using an auxin-inducible degron (AID) system leads to an increase in the overall size of the early endosome compartment [66]. We therefore tested whether *piki-1* mutants were able to suppress these defects. Consistent with previously reported results, we observed an overall increase in the size of the early endosome compartment in NEKL-2–depleted worms, as based on an ∼1.4-fold increase in mean intensity and size and an ∼1.2-fold increase in the number of puncta (Fig 4A, 4B, 4D–4F). In *piki-1(Q1507Stop)* mutants that were depleted of NEKL-2 relative to worms with NEKL-2 depletion alone, we observed a decrease in the mean intensity (∼1.3-fold), size (∼1.8-fold), and number of puncta (∼1.2-fold) (Fig 4B–4F). The early endosome compartment in NEKL-2–depleted *piki-1(Q1507Stop)* worms was comparable to the early endosome compartment in WT animals, albeit with a modest reduction in size (∼1.3-fold) (Fig 4A, 4C, 4D–4F). The suppression of the early endosome defects in *piki-1(Q1507Stop)* mutants at the early endosome indicates that PIKI-1 functions at the early endosome in *C. elegans* membrane trafficking.

**Fig 4.**
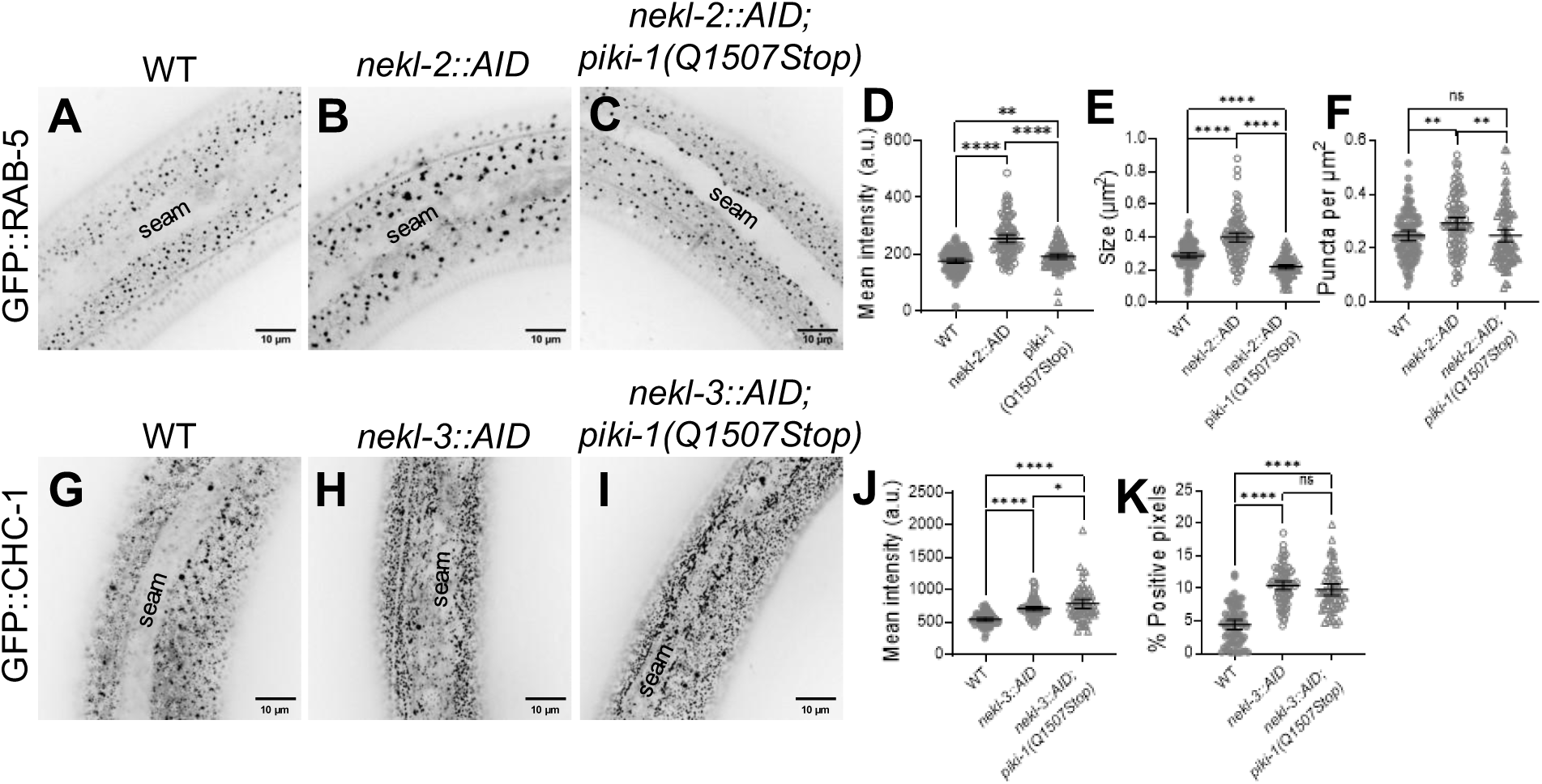
Loss of PIKI-1 suppresses *nekl-2*–associated defects in the early endosome. (A–C) Representative confocal images of P*_rab-5_*::GFP::RAB-5 (n = 96) expression in auxin-treated day 2 adults in (A) wild-type, (B) *nekl-2::AID*, and (C) *nekl-2::AID; piki-1(Q1507Stop)* backgrounds. (D–F) Dot plots show (D) mean intensity, (E) puncta size, and (F) the number of puncta per unit area for P*_rab-5_*::GFP::RAB-5 worms. (G–I) Representative confocal images of GFP::CHC-1 (n = 75) expression in auxin-treated day 2 adults in (G) wild-type, (H) *nekl-3::AID*, and (I) *nekl-3::AID; piki-1(Q1507Stop)* backgrounds. (J,K) Dot plots show the (I) mean intensity and (J) percent positive pixels (above threshold). (A–C, G–I) The seam cell is labeled in all images. (D–F,J, K) Dot plots show the mean and 95% CI. Statistical significance was determined by unpaired t-tests; ****p ≤ 0.0001, **p ≤ 0.01, *p ≤ 0.05; ns, not significant. Raw data are available in S1 File.

We showed above that PIKI-1 is associated with clathrin-coated pits, and strong defects in apical clathrin localization occur when NEKL-3 is depleted [65]. Given the colocalization of PIKI-1 with CLIC-1 and suppression of the RAB-5 defects, we hypothesized that PIKI-1 might also be able to suppress defects in clathrin. Consistent with prior reports, we observed a localization defect in GFP::CHC-1 when NEKL-3 is depleted, such that clathrin was concentrated at the apical surface of the epidermis. When compared with WT animals, NEKL-3–depleted worms showed an ∼1.3-fold increase in mean intensity, and NEKL-3–depleted *piki-1(Q1507Stop)* mutant worms showed an ∼1.4-fold increase in mean intensity (Fig 4G–4I). Moreover, with respect to the percentage of CHC-1::GFP–positive pixels (above threshold), there was an ∼2.4-fold increase in NEKL-3– depleted worms and an ∼2.2-fold increase in *piki-1(Q1507Stop)* mutants (p > 0.05; Fig 4G, 4H, 4J). These results indicate that loss of PIKI-1 is not able to suppress defects in clathrin caused by depletion of NEKL-3.

### Loss of PIKI-1 alters phosphoinositide pools in the epidermis

To understand how PIKI-1 as a Class II PI3K regulates membrane phospholipids and intersects with NEKL functions, we generated lipid sensors to investigate the distribution of phosphoinositide species in wild type and *piki-1* mutants. Class II PI3Ks contribute to the generation of two distinct lipid species, PI(3)P and PI(3,4)P_2_, which in turn recruit specific proteins to intracellular membrane compartments [14]. Expression of the GFP-based lipid biosensors was driven under the control of epidermal-specific promoters (P_hyp7_ or P*_nekl-3_*) [69,87,88], and each biosensor was integrated as a single copy using miniMos methods or was maintained as a stable extrachromosomal array (see S2 File) [78].

In mammalian cells, PI(3)P is associated with early endosomes as well as with autophagosomes [89]. Consistent with this, our PI(3)P sensor (P_hyp7_::2xFYVE::mNeonGreen) localized to punctate structures close to the apical surface in wild-type adults (Fig 5A, 5B). *piki-1(Q1507Stop)* mutants exhibited a similar pattern of PI(3)P localization, although we detected a slight decrease (∼1.2-fold) in the number of vesicles relative to wild type (Fig 5C). Furthermore, *piki-1(ok2346)* deletion mutants showed similar changes in PI(3)P localization (S4 Fig). The absence of a strong effect on PI(3)P is consistent with previous work demonstrating that class III PI3Ks are the predominant producers of PI(3)P [8,76]. To further support our findings, we carried out colocalization analysis with an early endosome marker, mKate2::RAB-5 [90]. Approximately 32% of the mKate2::RAB-5 signal overlapped with our PI(3)P sensor (Manders’ overlap = 0.32), and, as expected, ∼65% of PI(3)P-labeled puncta strongly overlapped with the mKate2:RAB-5 signal (Manders’ overlap = 0.65; Fig 6A–6D). This substantial overlap is further supported by the Pearson’s correlation coefficient (PCC = 0.37; Fig 6E). Thus, the PI(3)P pools that are slightly affected by loss of PIKI-1 are most likely associated with the early endosome. Moreover, our findings indicate that PI(3) is present on compartments other than RAB-5–marked early endosomes and that PI(3) localization may be confined to specific subdomains or subpopulations within early endosomes.

**Fig 5.**
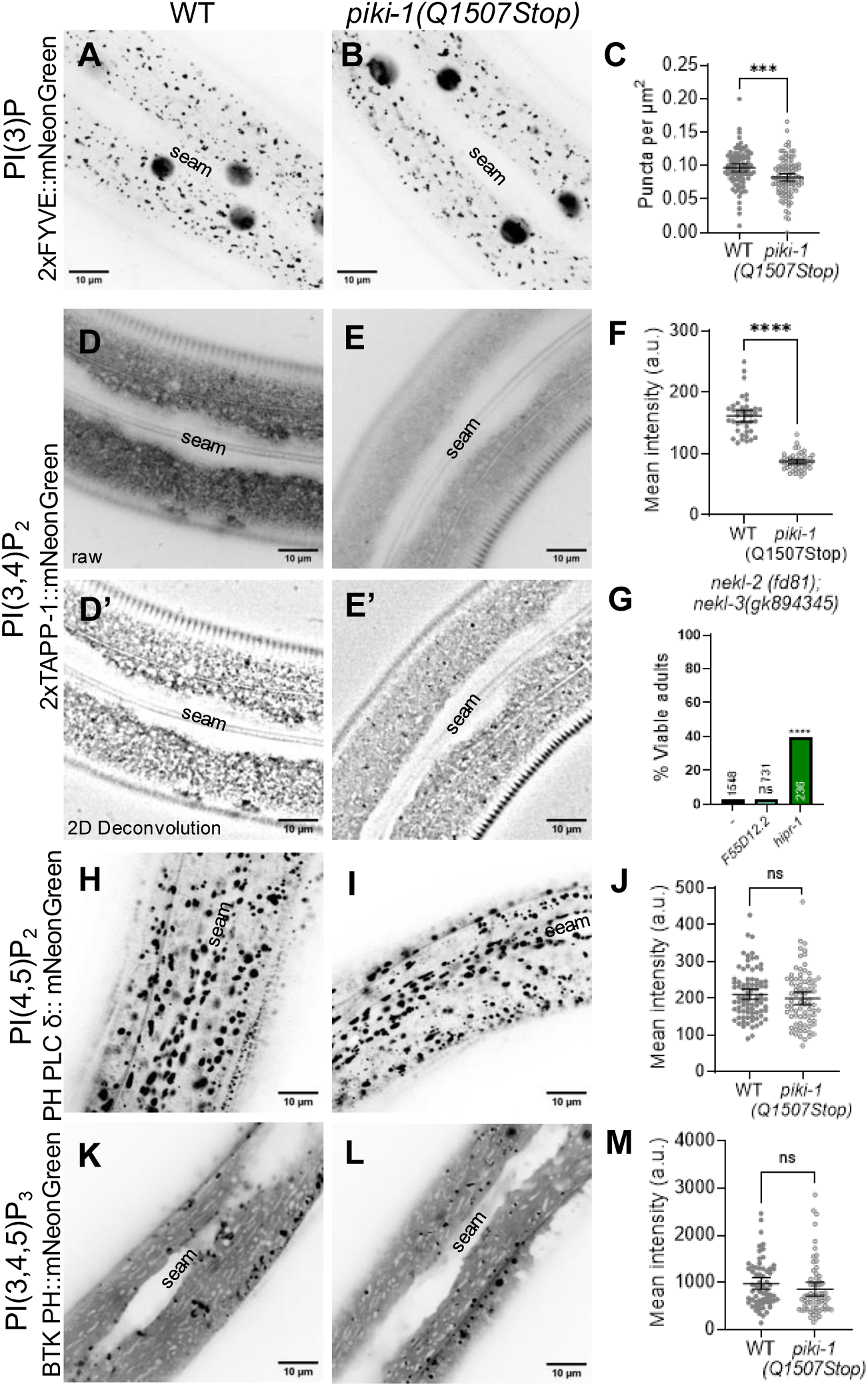
Loss of PIKI-1 affects PI(3)P and PI(3,4)P_2_ pools in the epidermis. (A,B,D,E,H,I,K,L) Representative confocal images of day-1 adult worms expressing (A, B) PI(3)P lipid sensor P_hyp7_::2xFYVE::mNeonGreen (n = 95), (D, E) PI(3,4)P_2_ lipid sensor P*_nekl-3_*::2xTAPP-1::mNeonGreen (n = 45), (H,I) PI(4,5)P_2_ lipid sensor P_hyp7_::PH PLC δ::mNeonGreen (n = 85), and (K,L) PI(3,4,5)P_3_ lipid biosensor P*_nekl-3_*::BTK PH::mNeonGreen (n = 64) in the (A,D,H,K) wild type and (B,E,I,L) *piki-1(Q1507Stop)* backgrounds. The seam cell is labeled in all images. (D’, E’) Representative confocal images of day-1 adult worms expressing P*_nekl-3_*::2xTAPP-1::mNeonGreen that have been processed using the 2D deconvolution algorithm in CellSens 4.2. (C) The puncta per unit area for worms expressing P_hyp7_::2xFYVE::mNeonGreen. (F,J,M) The mean intensity for worms expressing (F) P*_nekl-3_*::2xTAPP-1::mNeonGreen, (J) P_hyp7_::PH PLC δ::mNeonGreen, and (M) P*_nekl-3_*::BTK PH::mNeonGreen. Dot plots show the mean and 95% CI. (G) The percentage of viable adults after injection of dsRNA for *F55D12.2* and *hipr-1* relative to uninjected *nekl-2; nekl-3* worms. (C, F, G, J, M) Statistical significance was determined by unpaired t-tests; ****p ≤ 0.0001, ***p ≤ 0.001; ns, not significant. Raw data are available in S1 File.

**Fig 6.**
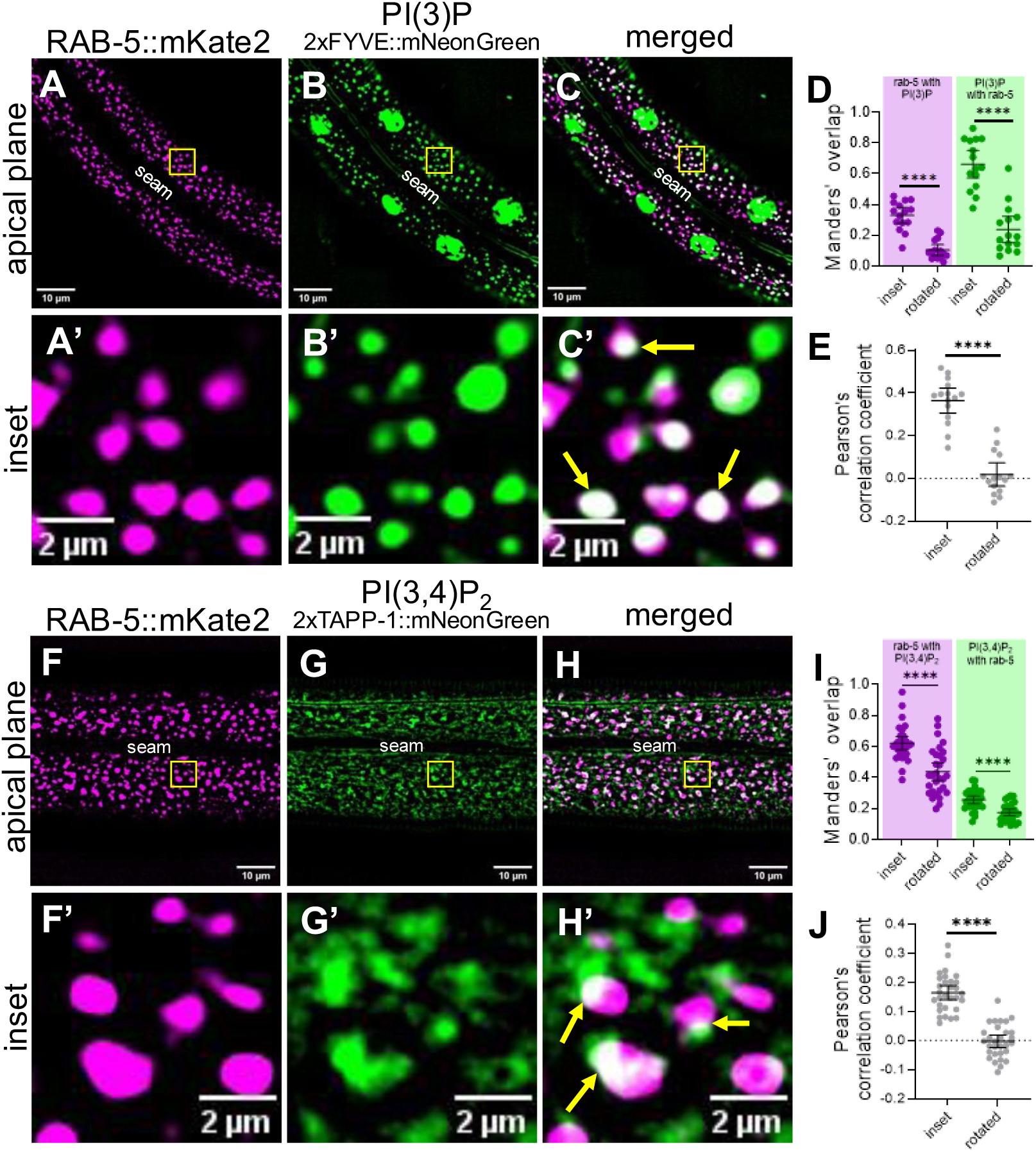
PI(3)P and PI(3,4)P_2_ localize at the early endosome. (A–C’, F−H’) Colocalization assays were carried out in worms expressing P*_dpy-7_*::mKate2::RAB-5 and (A–C’) the PI(3)P lipid sensor P_hyp7_::2xFYVE::mNeonGreen (n=15) and (F–H’) the PI(3,4)P_2_ lipid sensor P*_nekl-3_*::2xTAPP-1::mNeonGreen (n=30). The seam cell is labeled (A–H). Yellow squares (A–C, F–H) indicate enlarged insets (A’–C’, F’–H’). In merged insets (C’, H’), yellow arrows indicate examples of colocalization. (D, E, I, J) Colocalization was quantified using Mander’s overlap (D, I) and Pearson’s correlation coefficient (E, J). Dot plots show the mean and 95% CI. Statistical significance between rotated and inset values was determined by unpaired *t-* tests; ****p ≤ 0.0001. Raw data available in S1 File.

Like PI(3)P, PI(3,4)P is associated with early endosomes, as well as with endocytic recycling compartments [15,48]. Additionally, PI(3,4)P_2_ is found on the apical membrane of polarized epithelial cells [51]. In wild type, the PI(3,4)P_2_ sensor (P*_nekl-3_*::2xTAPP-1::mNeonGreen; *fdSi8*) exhibited a diffuse localization pattern in both apical and basal planes but also accumulated at apical puncta (Fig 6D, 6D’; S1 Movie). We note that the diffuse expression of the PI(3,4)P_2_ sensor may be due in part to the sensor being present in excess of its preferred ligand, a known challenge associated with genetically encoded lipid sensors [19]. Strikingly, reduction of PIKI-1 function led to a 1.9-fold decrease in the amount of the PI(3,4)P_2_ present in the epidermis (Fig 5D, 5D’, 5E, 5E’, 5F), consistent with PIKI-1 acting as a major producer of PI(3,4)P_2_. This decrease in the abundance of lipid-sensor positive structures may be attributable to a reduction in lipid-binding sites and increased reporter turnover, as has been reported [58–60,91,92]. Correspondingly, there was no observed accumulation of the lipid sensor at a more basal plane in *piki-1(Q1507Stop)* mutants (Movie).

Notably, our PI(3,4)P_2_ sensor (*fdEx406*) showed substantial colocalization with the mKate2::RAB-5 marker; ∼62% of the mKate2::RAB-5 signal overlapped with the PI(3,4)P_2_ sensor and ∼25% of the PI(3,4)P_2_ sensor overlapped with mKate2::RAB-5 (Fig 6F–6I). Significant positive but partial overlap was also supported by the Pearson’s correlation coefficient (PCC = 0.16; Fig 6J). These results suggest that a substantial proportion of early endosomes contain PI(3,4)P_2_, the production of which is dependent on PIKI-1. As with the PI(3)P sensor, our results suggest that PI(3,4)P_2_ may be associated with additional compartments, such as tubular elements emanating from early endosomes or apical recycling endosome compartments, and that PI(3,4)P_2_ may localize to specific subdomains of RAB-5–marked endosomes.

The phosphoinositide biosynthesis pathway is highly interconnected, with multiple lipid kinases and phosphatases contributing to the production of individual phosphoinositide species (Fig 1B). As such, changes in the levels of one phosphoinositide population have the potential to affect other pools [93]. To investigate whether loss of PIKI-1 affects other phosphoinositide pools, we generated lipid sensors to visualize PI(4,5)P_2_ (PH PLC δ::mNeonGreen), a phosphoinositide involved in clathrin-mediated endocytosis in mammalian cells, and PI(3,4,5)P_3_ (P*_nekl-3_*::BTK PH::mNeonGreen), a phosphoinositide directly upstream of PI(3,4)P_2_. The PI(4,5)P_2_ sensor, which localized throughout hyp7 to folded plasma membrane subdomain compartments that likely represent meisosomes [94], where PI(4,5)P_2_ is expressed,, appeared unchanged in *piki-1(Q1507Stop)* and *piki-1(ok2346)* worms (Fig 5H–5J; S4 Fig). Likewise, no obvious changes were observed for PI(3,4,5)P_3_ in *piki-1(Q1507Stop)* mutants, as this sensor showed a diffuse localization pattern but also accumulated at variably sized puncta throughout the epidermis (Fig 5K–5M). Our combined results indicate that reduction of PIKI-1 most strongly affects PI(3,4)P_2_ pools and has at most a limited impact on other species of phosphoinositides.

We also tested a putative dual-specificity lipid sensor reported to recognize both PI(3,4)P_2_ and PI(3,4,5)P_3_ (AKT::oxGFP) [59,95]. In wild-type animals, the localization of the dual-specificity sensor most closely resembled that of the PI(3,4)P_2_ sensor, with combined diffuse localization accompanied by distributed puncta (S5 Fig). Unexpectedly, the AKT::oxGFP reporter exhibited a dramatic tubulation phenotype in ∼50% of *piki-1(Q1507Stop)* and *piki-1(ok2346)* mutants (S5 Fig), reminiscent of the phenotype of *piki-1* mutants expressing TGN-38::GFP. Given that lysosomes have been reported to form tubules in the epidermis during molting cycles [96], we assessed two lysosomal markers, LMP-1::mNeonGreen [88] and NUC-1::mCherry [97], in *piki-1* mutants. LMP-1 is a conserved protein important for lysosome biogenesis [98] and NUC-1 is a lysosomal hydrolase [97]. Notably, we did not observe tubulation of either marker in *piki-1(Q1507Stop)* mutants (S6 Fig), indicating that the tubulations observed with the dual-specificity marker are unlikely to be affiliated with lysosomes. We note, however, that NUC-1::mCherry–positive vesicles were more abundant but slightly smaller in *piki-1(Q1507Stop)* mutants, suggesting that PIKI-1 may directly or indirectly affect lysosomal compartments (S6 Fig). Our findings suggest that the tubulations observed with the AKT::oxGFP marker, as well as with TGN-38::GFP, may be extensions of endosomal compartments that are not marked by any of our other tested markers. It is also possible that the AKT PH domain may retain the ability to bind other lipids or protein partners, thereby impacting its localization pattern.

### *nekl* defects may be suppressed by inhibition of PI(3,4)P_2_ binding partners

Our findings suggest that a reduction of PI(3,4)P_2_ is responsible for the genetic suppression of *nekl-2; nekl-3* mutants by *piki-1* mutants. Moreover, a reduction of PI(3,4)P_2_ would be expected to affect the binding of specific endocytic regulators to compartments including the early endosome. We therefore hypothesized that inhibition of one or more PI(3,4)P_2_-binding proteins may also lead to the suppression of *nekl-2; nekl-3* mutants. To test this, we looked for potential PI(3,4)P_2_-binding proteins involved in endocytic trafficking based on gene ontology (GO) terms. From this search, we identified two potential PI(3,4)P_2_ interactors: *F55D12.2*, an ortholog of human SESTD1, which regulates lipid signaling [99], and *hipr-1,* an ortholog of human HIP1 and HIPR1, proposed to regulate clathrin-mediated endocytosis and vesicular trafficking, by linking endocytic proteins to the actin cytoskeleton [100]. Notably, whereas inhibition of *F55D12.2* by RNAi failed to suppress molting defects in *nekl-2; nekl-3* worms, *hipr-1(RNAi)* resulted in ∼39% of *nekl-2; nekl-3* worms reaching adulthood (Fig 5G). Together, our findings support the model that reduced levels of PI(3,4)P_2_ on early endosomes, along with an accompanying reduction of PI(3,4)P_2_-binding proteins such as HIPR-1, lead to the suppression of *nekl-2; nekl-3* molting defects and to the partial restoration of trafficking functions.

## Discussion

Through a forward genetic screen aimed at identifying suppressors of *nekl-*associated molting defects, we isolated three independent alleles of the lipid modifier PIKI-1 and showed a novel connection between class II PI3K enzymes and NEKL kinases, which are essential for molting and membrane trafficking. Here we report that PIKI-1 regulates several endocytic processes, most notably through its role at early endosomes. PIKI-1 localizes to early endosomes, and its inhibition leads to a reduction in the size and number of vesicles marked with RAB-5 or EEA-1. Using genetically encoded lipid-biosensors [58,59], we found that PIKI-1 is a major contributor to membrane PI(3,4)P_2_ pools in the epidermis and that PI(3,4)P_2_ is found on the cytosolic leaflet of early endosomes. Our data support the model that class II PI3Ks, and not class I PI3Ks, are the predominant producers of PI(3,4)P_2_ in the *C. elegans* epidermis.

Reduction of PIKI-1 function also resulted in hyper-tubulation defects detected with markers for TGN-38, a trans-Golgi cargo that cycles through early endosomes on its way back to the Golgi; LGG-1, a marker for autophagosomes; and AKT-PH::GFP, a multi-specific lipid sensor. Although lysosomes in *C. elegans* form elongated tubules [96], our studies did not support a major role for PIKI-1 in controlling the morphology of the lysosomal compartment. Rather, we speculate that the observed tubulations may originate from early endosomes and that loss of PIKI-1 may lead to a defect in cargo sorting and/or endosomal tubule scission. The PI(3,4)P_2_ biosensor 2XTAPP1 appeared to label tubular or pleomorphic structures associated with RAB-5 positive puncta, suggesting the PI(3,4)P_2_ may be enriched on recycling tubules.

Colocalization studies indicated that PIKI-1 partially localizes to clathrin-coated pits as well as to early endosomes, consistent with findings in mammalian cells [10,12,14,38,48,70]. Studies in cell culture have indicated a role for class II PI3Ks in the production of PI(3,4)P_2_ at clathrin-coated pits, which then recruits proteins necessary for the completion of vesicle budding and the internalization of cargo [38,70]. Our findings, however, indicate that PIKI-1 is largely dispensable for clathrin-mediated endocytosis in the *C. elegans* epidermis. This discrepancy may reflect differences in the mechanisms controlling endocytosis in polarized cells versus non-polarized cells, as well as differences between cultured cells versus intact organisms [101].

Rather, our data support a major role for PIKI-1 in the production of PI(3,4)P_2_ at early endosomes and indicate that PIKI-1 has only a minor role in the production of PI(3), consistent with PI(3)P synthesis being driven principally by the class III PI3K VPS-34 [72,76]. Consistent with PIKI-1 being the major source of PI(3,4)P_2_, we failed to observe genetic interactions (suppression or enhancement) when other components of the phosphoinositide synthesis pathway were inhibited in *nekl-2; nekl-3* and *nekl-2; nekl-3 piki-1* backgrounds. These findings suggest that the conversion of PI(4)P to PI(3,4)P_2_ by PIKI-1 is the predominant means by which PI(3,4)P_2_ is produced in the epidermis. Our findings also support the model that inhibition of PIKI-1 leads to a reduction of PI(3,4)P_2_-binding proteins, which may ultimately impact NEKL functions. Consistent with this, inhibition of HIPR-1, the *C. elegans* ortholog of human HIPR1, a protein linking the endocytic machinery to the actin cytoskeleton, was sufficient to partially suppress *nekl-2; nekl-3* molting defects.

Previous studies in *C. elegans* embryos and in the adult gonad indicate that PIKI-1 has a primary role in autophagy in these contexts [72,74–76]. For example, PI(3)P production, which controls autophagy in the *C. elegans* embryo, is dependent on the coordination among PIKI-1, VPS-34, and MTM-1 (a 3-phosphatase). In addition, loss of PIKI-1 in the embryo and in the gonad strongly impairs autophagosome clearance [76]. By contrast, reduction of PIKI-1 function had a relatively minor impact on autophagosomes in the epidermis (marked by mNeonGreen::LGG-1), leading to a decrease in marker intensity and a modest increase in tubulation. These data suggest that the role of PIKI-1 in *C. elegans* is at least partially cell-type specific. Cell-type specificity of class II PI3K activity in human cells may be accomplished, in part, through the expression of different PI3K isoforms in different tissues [8,10].

In previous studies of the *nekls*, we identified genetic suppressors with well-established functions in membrane trafficking including proteins controlling clathrin-mediated endocytosis, membrane lipid asymmetry, and endosomal-associated actin [65–69,102]. Other suppressors act independently of membrane trafficking and include regulators of cargo processing, cell signaling, and development [103–105]. Our findings indicate that loss of PIKI-1 leads to *nekl* suppression through a mechanism that does affect membrane trafficking. Importantly, reduction of PIKI-1 activity at least partially corrected defects in early endosomes caused by loss of NEKL-2, a finding that dovetails with our results showing that PIKI-1 acted at early endosomes. Our results suggest that PIKI-1 and NEKL-2 may act in opposition and that PIKI-1 could be a direct or indirect target for negative regulation by the NEKLs. Interestingly, we recently identified PIKI-1 as a proximal interactor of NEKL-2 and NEKL-3 in the epidermis, raising the possibility that PIKI-1 may be directly regulated by the NEKLs. Future studies combining genetic, proteomic, and biochemical approaches are expected to provide novel insights into how conserved NEKL kinases regulate multiple steps of endocytic trafficking.

## Materials and Methods

### Strains and propagation

All *C. elegans* strains were maintained per standard protocols and propagated at 22°C unless stated otherwise [106]. Strains in this study are listed in S1 Table in S2 File.

### RNAi

Standard dsRNA injection methods were used to conduct RNAi experiments [107]. Primers containing the T7 RNA polymerase–binding motif and corresponding to *piki-1, age-1, inpp-1, inpp-4b, F55D12.2,* and *hipr-1* were used to synthesize dsRNA using the MEGAscript RNAi Kit (Invitrogen). dsRNA was injected at concentrations of 500–1000 ng/μL. Primer information is in S2 Table in S2 File.

### CRISPR mutant alleles

Alleles of *piki-1* (Q1507Stop) were created using established CRISPR-Cas9 protocols [108–110]. sgRNA and repair templates were synthesized by Integrated DNA technologies and Dharmacon-Horizon Discovery; ApE and CRISPRcruncher were used in the design of the guideRNA and repair templates [111,112]. Primer, sgRNA, and repair template sequences are provided in S3 Table in S2 File.

### Reporter strain construction

Plasmids for *C. elegans* epidermal-specific expression in the hyp7 syncytium used promoters for *semo-1*/Y37A1B.5 (P_hyp7_) and *nekl-3* (P*_nekl-3_*). We generated pDONR221 entry vectors containing coding regions for *C. elegans piki-1* (gift from Zheng Zhou, Baylor College of Medicine) [76], human *2x-TAPP-1,* wherein the second repeat was codon optimized for *C. elegans* [113] and contained synthetic introns to promote expression (Integrated DNA Technologies); human *AKT* [95]; and human *BTK* (Addgene Plasmid #51463) [114]. Cloning of the PH-domain from human AKT (pDONR221 AKT) [95]into destination vector pCFJ1662 P*_semo-1_*::GTWY::oxGFP::let-858 (35G7) [66,87,88]was performed using the Gateway LR clonase II reaction (Invitrogen). *piki-1* and *BTK* pDONR221 clones were transferred into destination vector pCFJ1662 P_hyp7_::GTWY::mNeonGreen::let-858 (34H4) [88] via the Gateway LR clonase II reaction. The *2X-TAPP-1* pDONR 221 clone was transferred into destination vector P*_nekl-3_::*GTWY::mNeonGreen::let-858 (pDF477, derived from pCFJ1662 (34H4)) [69] using the Gateway LR clonase II reaction. To generate single-copy integrations, standard miniMos procedures were followed [78].

In cases where we were unable to obtain integrations, we co-injected the expression clone and a plasmid containing *unc-119(+)* into a background of *unc-119(ed3)* and used the resulting stable extrachromosomal array in our analyses (Fig 5K, 5L, 6G).

### Auxin treatment

Indole-3-acetic acid (auxin) from Alfa Aesear was used to make a 100× stock auxin solution (0.4 M) by dissolving 0.7 g of auxin in 10 mL of 100% ethanol. For experiments, a mixture of 25 µL of stock auxin solution and 225 µL of autoclaved deionized water was added to NGM plates spotted with OP50 with day-1 adult worms present, at least 18 hours before imaging [66,115].

### Image acquisition

Fluorescence images were acquired using an Olympus IX83 P2ZF inverted microscope with a Yokogawa spinning-disc confocal head (CSU-W1). z-Stack images were acquired using a 100×1.35 N.A. silicone oil objective. cellSens 4.2 software (Olympus Corporation) was used for image acquisition with a Hamamatsu-ORCA-Fusion camera. For each worm, z-stack slices were acquired every 0.2 µm for ∼20 slices to encompass the epidermis of the worm.

### Image analysis to determine fluorescence, size, puncta per unit area, and percent positive pixels

All image analysis and quantification, was done using Fiji [116].

To quantify the mean intensity (measured in arbitrary units, a.u.), the intensity of the background of the image was first measured using the rectangle selection tool in an area of the image where there was no visible fluorescence. The resulting value was subtracted from the mean intensity value obtained from the epidermis (hyp7) of each animal in each picture, by using the polygon selection tool to select the appropriate region of interest (ROI) [66,69].

To quantify the average area of vesicles, number of puncta, or the percent positive pixels above threshold for a z-plane of interest which corresponds to a single slice at an appropriate plane from the obtained z-stacks, images underwent processing to remove background. This was done through application of the rolling ball background subtraction method (in which background intensity values below the average within a 50-pixel radius surrounding a positive pixel are subtracted; Fig 3F, 3G, 3N–3Q, S3 Fig; S6 Fig). For images for which rolling ball background subtraction did not work well, we used the minimum filter method (in which the central pixel is compared to other pixels within a 10-pixel radius and the minimum value in the window is replaced with the central pixel value to reduce noise; Fig 3B, 3C, 3H, 3I; S3 Fig). The filtered image was subtracted from the raw image using the image calculator function. After processing, all images were thresholded using the algorithm that worked best (or a manual threshold set using representative images; see S1 File for details and raw data). The “Despeckle” function was subsequently applied to all images to remove signal noise of ≤1pixel in size. Then the “Analyze Particles” function was applied to the processed images to determine the average area of vesicles. Within each experiment, the same background subtraction and threshold algorithms were used for all images. To account for variation in the size of the region of interest (ROI; drawn using the polygon tool to select hyp7 and exclude the seam cell) measured among worms, the number of puncta was divided by the area of the ROI (output in units of puncta per square micron).

### Image analysis to determine colocalization

For quantifying colocalization, the raw z-stack images were deconvoluted using the 2D deconvolution algorithm available in cellSens (ver. 4.2). The appropriate z-plane was then extracted from both the raw and deconvoluted images for each channel. To obtain a binary image to be used as a mask, deconvoluted images were thresholded. This binary mask was then combined with the raw image using the “AND” Boolean operation. The ROI was drawn around hyp7, excluding the seam cell, by using the polygon tool. To calculate the Pearson’s correlation coefficient (R) and Mander’s overlap (M) for these experiments, we used the BIOP JACoP plugin [117]. Merged images (containing both red and green channels) were used to determine the cases of significant overlap versus random co-occurrence. From these merged images, a small inset of 100 × 100 pixels (10,000 pixels^2^) was sampled, and the R and M values were calculated using the BIOP JACoP plugin (shown as “inset” in the resulting graphs). To create a random distribution of green and red pixels of interest as a control against random coincidence, the red channel was rotated 90° in relation to the green channel using the transform function before the R and M values were calculated using the BIOP JACoP plugin (shown as “rotated”) [69,118].

For colocalization between PIKI-1::mNeonGreen and mKate2::RAB-5, we generated trans-heterozygous worms. Worms homozygous for PIKI-1::mNeonGreen and mKate2::RAB-5 exhibited ectopic expression of PIKI-1::mNeonGreen in the seam cell and had almost no expression in hyp7 whereas mKate2::RAB-5 expression appeared unchanged (S7 Fig). This was done by crossing N2 males to hermaphrodites expressing mKate2::RAB-5. The resulting red F1 males were subsequently crossed with hermaphrodites expressing PIKI-1::mNeonGreen. Red L4s were picked ∼18 hours before imaging as day-1 adults.

### Image analysis for phenotypes

Worms expressing P_hyp7_::AKT::GFP were scored based on the appearance of the marker throughout the entire epidermis. Worms were considered to have diffuse expression if there was a uniform architecture of the GFP signal and the boundary between hyp7 and the seam cell was clear. Worms were considered to have aggregates if there were at least three large aggregations present within the z-stack that were not uniform in size or morphology and were distinct from the background and nuclear expression. Worms with tubulations had elongated compartments originating from two or more sites in hyp7. All worms expressing AKT::GFP that exhibited tubulations also contained aggregations.

Worms expressing TGN-38::GFP or mNeonGreen::LGG-1 were scored based on the absence or presence of tubulations within the epidermis. Tubulations were defined as being longer, extended vesicles or longer, thin protrusions extending through the epidermis.

### Statistical analysis

GraphPad Prism software was used to perform statistical tests in accordance with standard methods [119].

## Acknowledgements

The authors thank Amy Fluet for editing this manuscript. The authors also thank Zheng Zhou (Baylor College of Medicine) for the plasmid construct containing *piki-1* cDNA and Reto Gassmann (University of Porto) for sharing GCP1474 (*P_dpy7_::mKate-2::RAB-5)* with us. Some strains were provided by the CGC, which was funded by the NIH Office of Research Infrastructure Programs (P40 OD010440). The authors acknowledge the Center for Advanced Scientific Instrumentation (CASI) at the University of Wyoming for access to spINBRE TIRF. This research was supported by an Institutional Development Award (IDeA) from the National Institute of General Medical Sciences of the NIH under Grant #2P20GM103432.

## Supporting Information

**S1 Fig.**
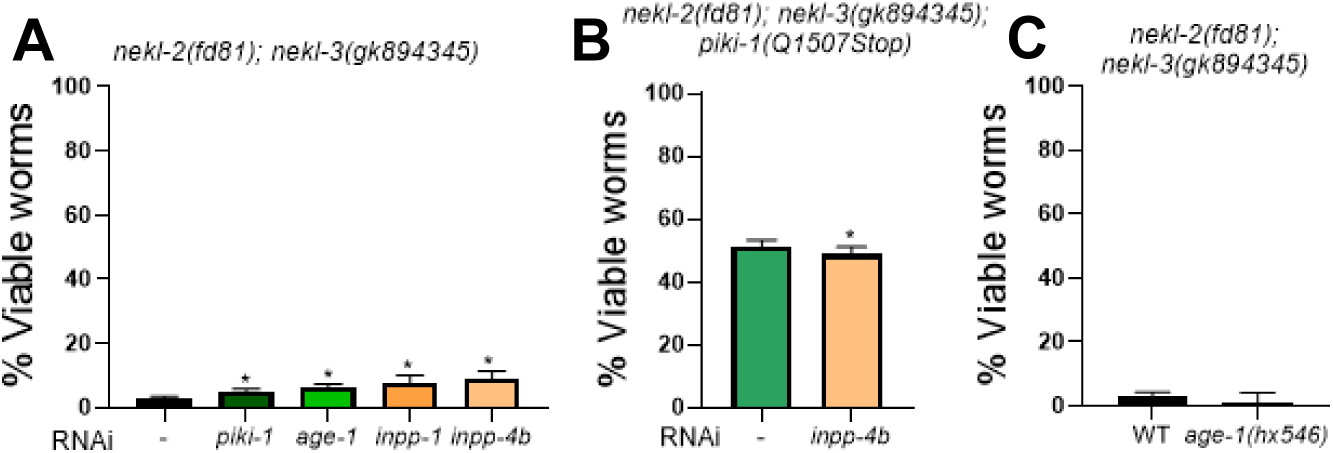
Supplemental RNAi and suppression data. (A) The proportion of viable progeny produced by *nekl-2(fd81); nekl-3(gk894345)* worms after injection by the indicated dsRNA. (B) The proportion of viable progeny produced by *nekl-2(fd81); nekl-3(gk894345) piki-1(Q1507Stop)* worms after injection by the indicated dsRNA. (C) The proportion of viable progeny produced by *nekl-2(fd81); nekl-3(gk894345)* worms for the indicated genotype. Statistical significance was determined using unpaired t-tests; *p≤0.05. Raw data available in S1 File.

**S2 Fig.**
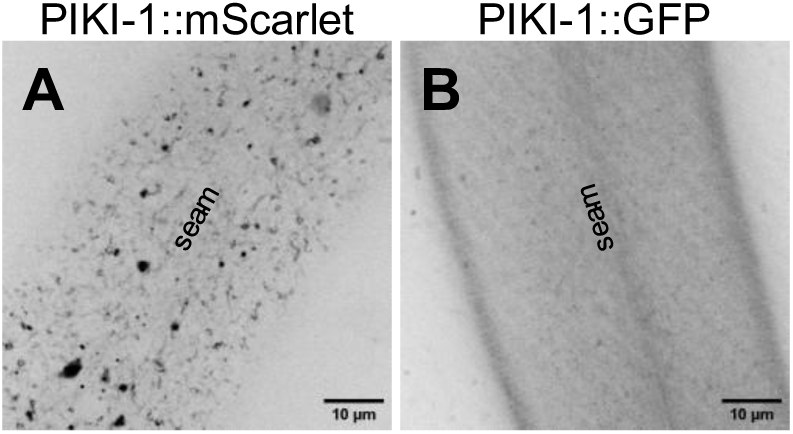
Endogenous PIKI-1 expression. (A, B) Representative confocal images of day-1 adults expressing (A) PIKI-1::mScarlet and (B) PIKI-1::GFP.

**S3 Fig.**
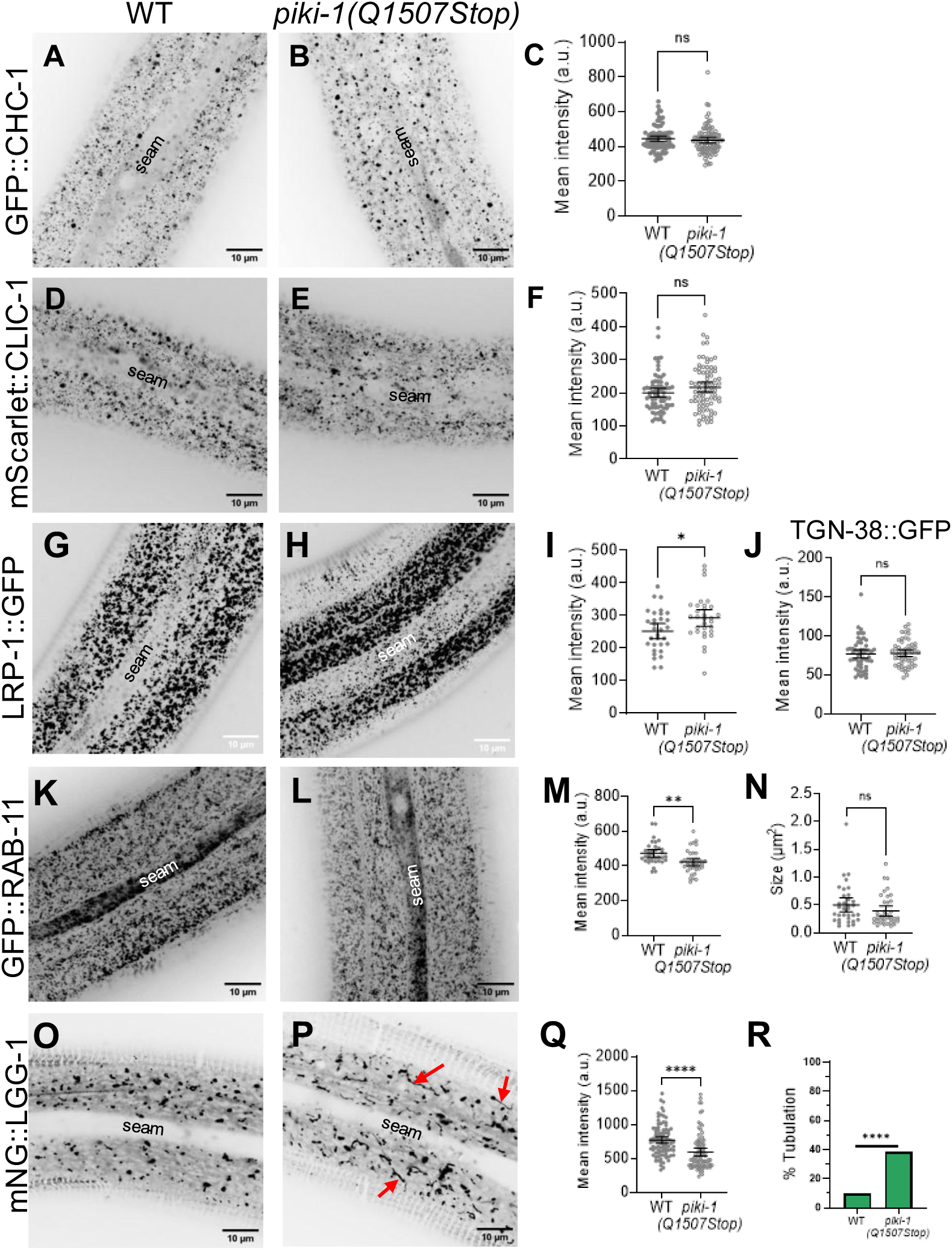
Effects of reduction of PIKI-1 function on intracellular trafficking compartments and cargo. (A,B,D,E,G,H,K,L,O,P) Representative confocal microscopy images of day-1 adults to assess the effects of *piki-1(Q1507Stop)* mutants relative to wild-type worms with respect to (A,B) GFP::CHC-1 (n=93), (D,E) mScarlet::CLIC-1 (n=79), (G,H) P_hyp7_::LRP-1::GFP (n=31), (K,L) P_hyp7_::GFP::RAB-11, and (O,P) P_hyp7_::mNeonGreen::LGG-1. Red arrows in (P) indicate instances of tubulation. The seam cell is labeled in all images. (C, F, I, M, Q) The mean intensity for each marker tested in (A,B) GFP::CHC-1, (D,E) mScarlet::CLIC-1, P_hyp7_::LRP-1::GFP, (K,L) P_hyp7_::GFP::RAB-11, and (O,P) P_hyp7_::mNeonGreen::LGG-1. (J) The mean intensity for P_hyp7_::TGN-38::GFP. (N) The size of vesicles was graphed onto a dot plot for the P_hyp7_::GFP::RAB-11 strains. (R) The percentage of worms that had either no tubulations or tubulations was recorded for P_hyp7_::mNeonGreen::LGG-1. (C, F, I, J, M, N, Q) Dot plots show the mean and 95% CI. Statistical significance was determined by unpaired *t-*tests; ****p ≤ 0.0001, **p ≤ 0.01, *p ≤ 0.05; ns, not significant. (R) Statistical significance was determined by Fisher’s exact test; ****p ≤ 0.0001. Raw data are available in S1 File.

**S4 Fig.**
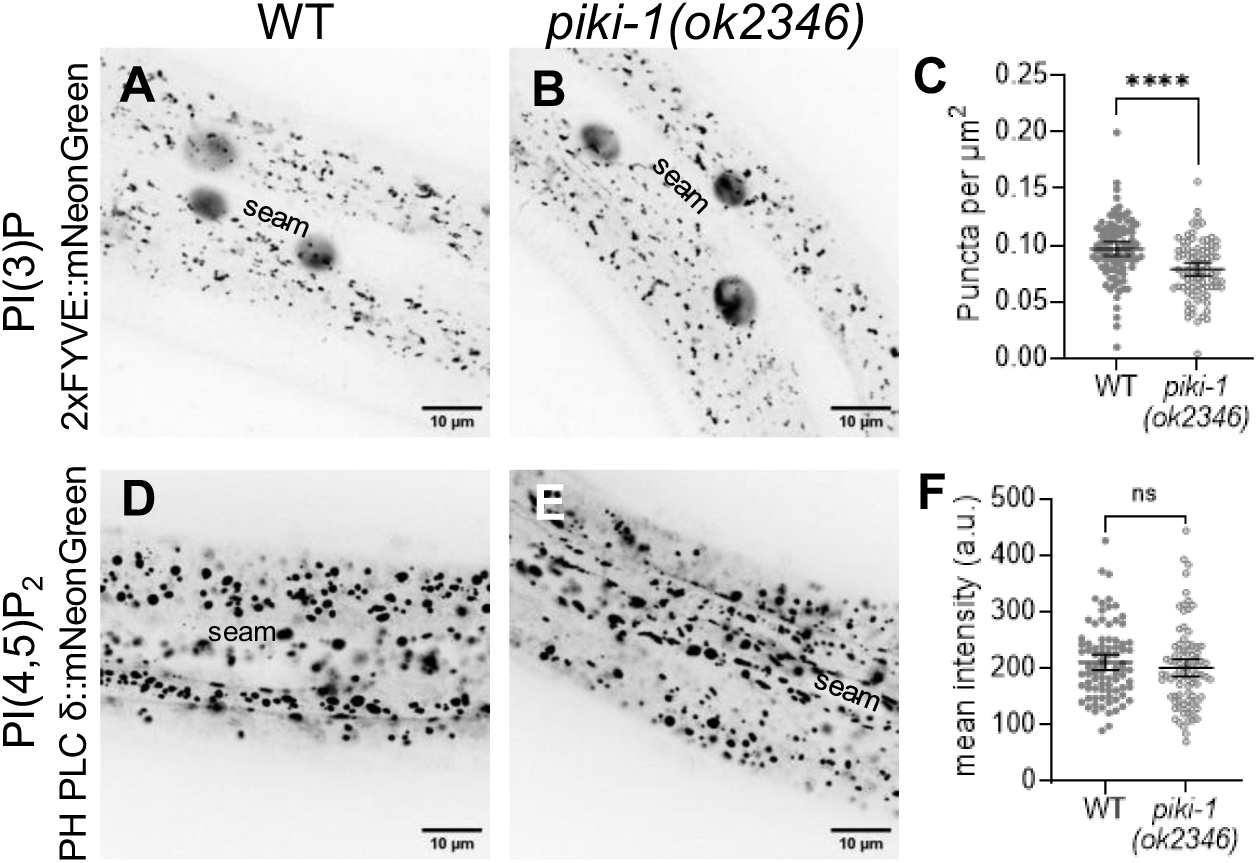
Effects of *piki-1(ok2346)* deletion allele on PI(3)P and PI(4,5)P_2_. (A,B,D,E) Representative confocal images of day-1 adults show the effects in *piki-1(ok2346)* mutants relative to wild-type worms with respect to (A, B) the PI(3)P lipid sensor P_hyp7_::2xFYVE::mNeonGreen and (D,E) the PI(4,5)P_2_ lipid sensor P_hyp7_::PH PLC δ::mNeonGreen. (C) Puncta per unit area for worms expressing P_hyp7_::2xFYVE::mNeonGreen. (F) Mean intensity for worms expressing P_hyp7_::PH PLC δ::mNeonGreen. Dot plots show the mean and 95% CI. Statistical significance was determined by unpaired *t-*tests; *p ≤ 0.05; ns, not significant. Raw data are available in S1 File.

**S5 Fig.**
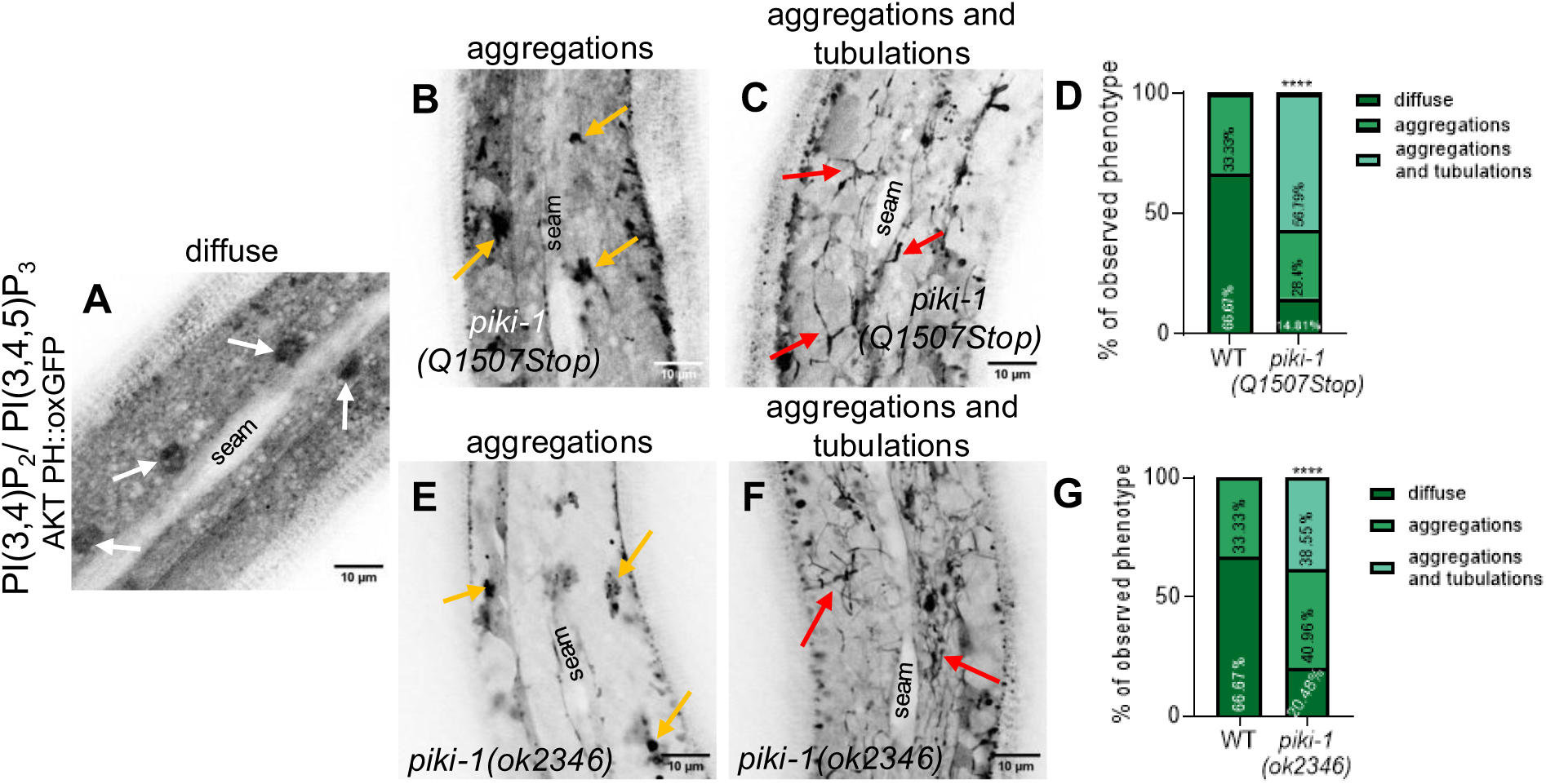
Effects of reduction of PIKI-1 function on the multi-specific lipid sensor AKT-PH. (A–C,E,F) Representative confocal images of day-1 adults that expressed the PI(3,4)P_2_/PI(3,4,5)P_3_ lipid sensor P_hyp7_::AKT::oxGFP in the (A) wild-type, (B,C) *piki-1(Q1507Stop)* (n=81), or (E,F) *piki-1(ok2346)* (n = 83) background. White arrows (A) indicate nuclei. Gold arrows (B, E) indicate aggregations. Red arrows (C, F) indicate tubulations. (D, G) Worms expressing P_hyp7_::AKT-PH::oxGFP were scored for the presence of diffuse labeling, of aggregations, and of aggregations and tubulations within the epidermis. Statistical significance of the differences in phenotype distribution across backgroundswas determined by Fisher’s exact test; ****p < 0.0001. Raw data are available in S1 File.

**S6 Fig.**
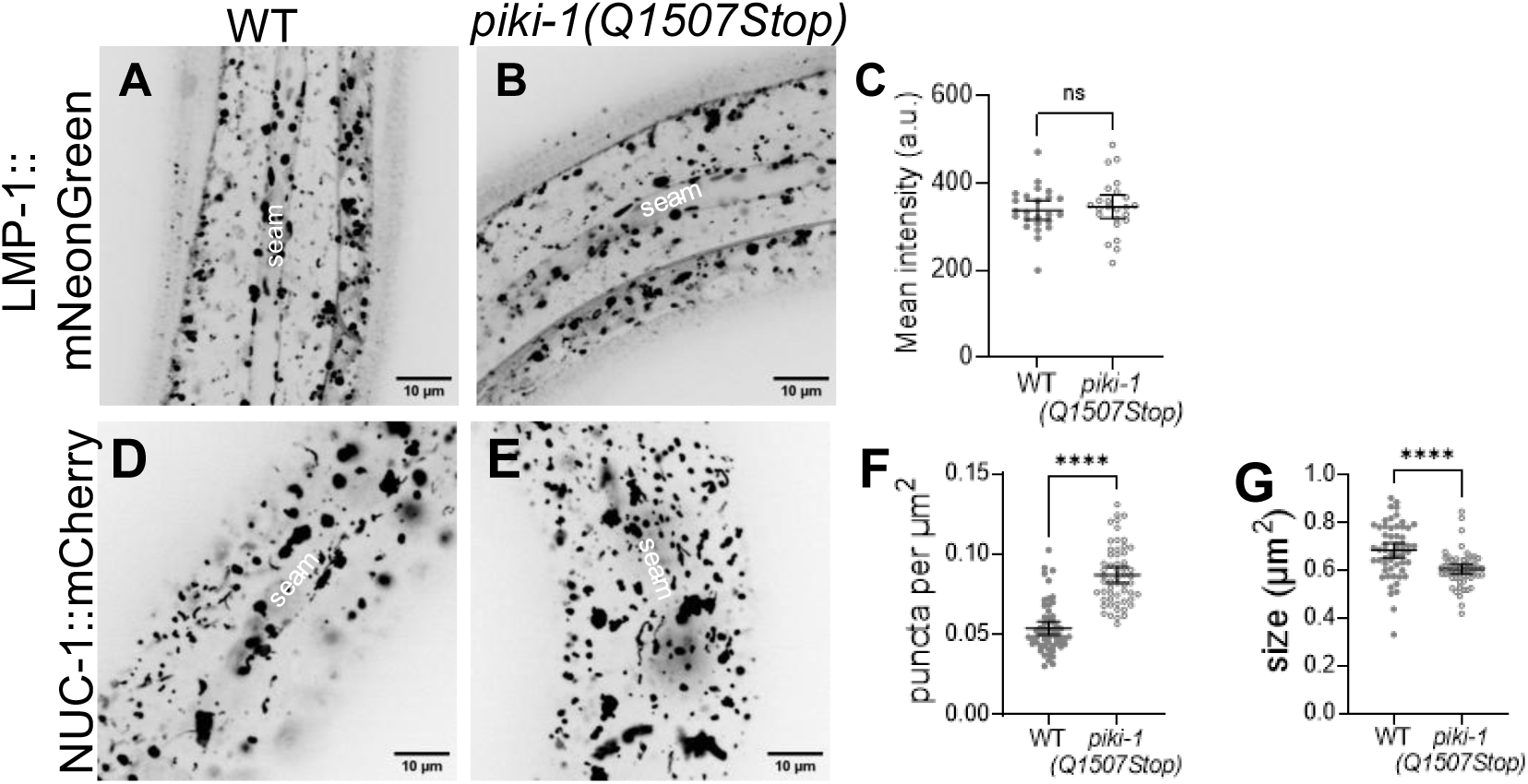
Effects of reduction of PIKI-1 function on lysosomes. (A,B,D,E) Representative confocal images of day-1 adults expressing (A,B) P_hyp7_::LMP-1::mNeonGreen (n=24) or (D,E) NUC-1::mCherry (n=56) in the (A,D) wild-type and (B,E) *piki-1(Q1507Stop)* backgrounds. (C) Mean intensity was plotted for P_hyp7_::LMP-1::mNeonGreen. (F,G) The (F) number of puncta per unit area and (G) size were plotted for NUC-1::mCherry. Dot plots show the mean and 95% CI. Statistical significance was determined by unpaired *t-*tests; ****p ≤ 0.0001; ns, not significant.

**S7 Fig.**
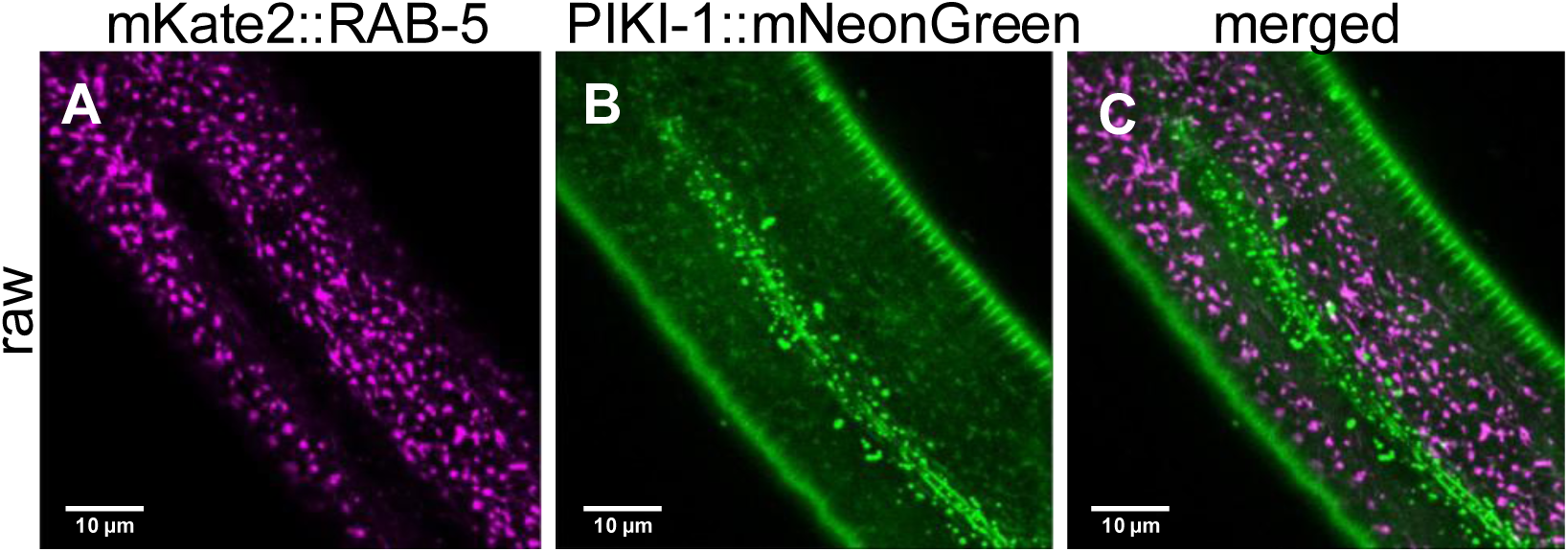
Marker expression in worms homozygous for P_hyp7_::PIKI-1::mNeonGreen and P*_dyp-7_*::mKate2::RAB-5. (A–C) Representative raw images of a day-1 adult homozygous for both P_hyp7_::PIKI-1::mNeonGreen and P*_dpy-7_*::mKate2::RAB-5. Both (A,B) single-channel and (C) merged images are shown.

**S1 Movie. P*_nekl-3_*::2xTAPP-1::mNeonGreen in the epidermis of a wild-type worm.**

z-Stack movie showing the expression of the PI(3,4)P_2_ sensor P*_nekl-3_*::2xTAPP-1::mNeonGreen from the apical to the basal plane in the epidermis of a wild-type worm.

**S2 Movie. P*_nekl-3_*::2xTAPP-1::mNeonGreen in the epidermis of a *piki-1(Q1507Stop)* worm.**

z-Stack movie showing the expression of the PI(3,4)P_2_ sensor P*_nekl-3_*::2xTAPP-1::mNeonGreen from the apical to the basal plane in the epidermis of a *piki-1(Q1507Stop)* mutant.

## References

1. Dickson EJ, Hille B. Understanding phosphoinositides: rare, dynamic, and essential membrane phospholipids. Biochem J. 2019;476: 1–23. doi:10.1042/BCJ20180022

2. Balla T. Phosphoinositides: Tiny Lipids With Giant Impact on Cell Regulation. Physiol Rev. 2013;93: 1019–1137. doi:10.1152/physrev.00028.2012

3. Falkenburger BH, Jensen JB, Dickson EJ, Suh B-C, Hille B. Phosphoinositides: lipid regulators of membrane proteins. J Physiol. 2010;588: 3179–3185. doi:10.1113/jphysiol.2010.192153

4. Haucke V, Di Paolo G. Lipids and lipid modifications in the regulation of membrane traffic. Curr Opin Cell Biol. 2007;19: 426–435. doi:10.1016/j.ceb.2007.06.003

5. Posor Y, Jang W, Haucke V. Phosphoinositides as membrane organizers. Nat Rev Mol Cell Biol. 2022;23: 797–816. doi:10.1038/s41580-022-00490-x

6. Marat AL, Haucke V. Phosphatidylinositol 3-phosphates—at the interface between cell signalling and membrane traffic. EMBO J. 2016;35: 561–579. doi:10.15252/embj.201593564

7. Di Paolo G, De Camilli P. Phosphoinositides in cell regulation and membrane dynamics. Nature. 2006;443: 651–657. doi:10.1038/nature05185

8. Bilanges B, Posor Y, Vanhaesebroeck B. PI3K isoforms in cell signalling and vesicle trafficking. Nat Rev Mol Cell Biol. 2019;20: 515–534. doi:10.1038/s41580-019-0129-z

9. Burke JE, Triscott J, Emerling BM, Hammond GRV. Beyond PI3Ks: targeting phosphoinositide kinases in disease. Nat Rev Drug Discov. 2023;22: 357–386. doi:10.1038/s41573-022-00582-5

10. Jean S, Kiger AA. Classes of phosphoinositide 3-kinases at a glance. J Cell Sci. 2014;127: 923–928. doi:10.1242/jcs.093773

11. Walker EH, Perisic O, Ried C, Stephens L, Williams RL. Structural insights into phosphoinositide 3-kinase catalysis and signalling. Nature. 1999;402: 313–320. doi:10.1038/46319

12. Safaroghli-Azar A, Sanaei M-J, Pourbagheri-Sigaroodi A, Bashash D. Phosphoinositide 3-kinase (PI3K) classes: From cell signaling to endocytic recycling and autophagy. Eur J Pharmacol. 2023;953: 175827. doi:10.1016/j.ejphar.2023.175827

13. Thibault B, Ramos-Delgado F, Guillermet-Guibert J. Targeting Class I-II-III PI3Ks in Cancer Therapy: Recent Advances in Tumor Biology and Preclinical Research. Cancers. 2023;15: 784. doi:10.3390/cancers15030784

14. Yoshioka K. Class II phosphatidylinositol 3-kinase isoforms in vesicular trafficking. Biochem Soc Trans. 2021;49: 893–901. doi:10.1042/BST20200835

15. Hawkins P, Stephens L. Emerging evidence of signalling roles for PI(3,4)P2 in Class I and II PI3K-regulated pathways. Biochem Soc Trans. 2016;44: 307–314. doi:10.1042/BST20150248

16. Gulluni F, Santis MCD, Margaria JP, Martini M, Hirsch E. Class II PI3K Functions in Cell Biology and Disease. Trends Cell Biol. 2019;29: 339–359. doi:10.1016/j.tcb.2019.01.001

17. Marcelić M, Mahmutefendić Lučin H, Jurak Begonja A, Blagojević Zagorac G, Lučin P. Early Endosomal Vps34-Derived Phosphatidylinositol-3-Phosphate Is Indispensable for the Biogenesis of the Endosomal Recycling Compartment. Cells. 2022;11: 962. doi:10.3390/cells11060962

18. Roggo L, Bernard V, Kovacs AL, Rose AM, Savoy F, Zetka M, et al. Membrane transport in Caenorhabditis elegans: an essential role for VPS34 at the nuclear membrane. EMBO J. 2002;21: 1673–1683. doi:10.1093/emboj/21.7.1673

19. Goulden BD, Pacheco J, Dull A, Zewe JP, Deiters A, Hammond GRV. A high-avidity biosensor reveals plasma membrane PI(3,4)P2 is predominantly a class I PI3K signaling product. J Cell Biol. 2018;218: 1066–1079. doi:10.1083/jcb.201809026

20. Gozzelino L, De Santis MC, Gulluni F, Hirsch E, Martini M. PI(3,4)P2 Signaling in Cancer and Metabolism. Front Oncol. 2020;10. Available: https://www.frontiersin.org/articles/10.3389/fonc.2020.00360

21. Wang H, Loerke D, Bruns C, Müller R, Koch P-A, Puchkov D, et al. Phosphatidylinositol 3,4-bisphosphate synthesis and turnover are spatially segregated in the endocytic pathway. J Biol Chem. 2020;295: 1091–1104. doi:10.1074/jbc.RA119.011774

22. Yudushkin I. TAPPing into PIPs: A new reporter reveals the origin of plasma membrane PI(3,4)P2. J Cell Biol. 2019;218: 735–736. doi:10.1083/jcb.201901143

23. Stephens LR, Jackson TR, Hawkins PT. Agonist-stimulated synthesis of phosphatidylinositol(3,4,5)-trisphosphate: A new intracellular signalling system? Biochim Biophys Acta BBA - Mol Cell Res. 1993;1179: 27–75. doi:10.1016/0167-4889(93)90072-W

24. Watt SA, Kimber WA, Fleming IN, Leslie NR, Downes CP, Lucocq JM. Detection of novel intracellular agonist responsive pools of phosphatidylinositol 3,4-bisphosphate using the TAPP1 pleckstrin homology domain in immunoelectron microscopy. Biochem J. 2004;377: 653–663. doi:10.1042/bj20031397

25. Koch PA, Dornan GL, Hessenberger M, Haucke V. The molecular mechanisms mediating class II PI 3-kinase function in cell physiology. FEBS J. 2021;288: 7025–7042. doi:10.1111/febs.15692

26. Lo W-T, Zhang Y, Vadas O, Roske Y, Gulluni F, De Santis MC, et al. Structural basis of phosphatidylinositol 3-kinase C2α function. Nat Struct Mol Biol. 2022;29: 218–228. doi:10.1038/s41594-022-00730-w

27. Lo W-T, Belabed H, Kücükdisli M, Metag J, Roske Y, Prokofeva P, et al. Development of selective inhibitors of phosphatidylinositol 3-kinase C2α. Nat Chem Biol. 2023;19: 18–27. doi:10.1038/s41589-022-01118-z

28. Misawa H, Ohtsubo M, Copeland NG, Gilbert DJ, Jenkins NA, Yoshimura A. Cloning and Characterization of a Novel Class II Phosphoinositide 3-Kinase Containing C2 Domain. Biochem Biophys Res Commun. 1998;244: 531–539. doi:10.1006/bbrc.1998.8294

29. Wang H, Lo W-T, Žagar AV, Gulluni F, Lehmann M, Scapozza L, et al. Autoregulation of Class II Alpha PI3K Activity by Its Lipid-Binding PX-C2 Domain Module. Mol Cell. 2018;71: 343–351.e4. doi:10.1016/j.molcel.2018.06.042

30. Wheeler M, Domin J. The N-terminus of phosphoinositide 3-kinase-C2β regulates lipid kinase activity and binding to clathrin. J Cell Physiol. 2006;206: 586–593. doi:10.1002/jcp.20507

31. Redpath GMI, Betzler VM, Rossatti P, Rossy J. Membrane Heterogeneity Controls Cellular Endocytic Trafficking. Front Cell Dev Biol. 2020;8. doi:10.3389/fcell.2020.00757

32. Gaidarov I, Smith MEK, Domin J, Keen JH. The Class II Phosphoinositide 3-Kinase C2α Is Activated by Clathrin and Regulates Clathrin-Mediated Membrane Trafficking. Mol Cell. 2001;7: 443–449. doi:10.1016/S1097-2765(01)00191-5

33. Daste F, Walrant A, Holst MR, Gadsby JR, Mason J, Lee J-E, et al. Control of actin polymerization via the coincidence of phosphoinositides and high membrane curvature. J Cell Biol. 2017;216: 3745–3765. doi:10.1083/jcb.201704061

34. Höning S, Ricotta D, Krauss M, Späte K, Spolaore B, Motley A, et al. Phosphatidylinositol-(4,5)-Bisphosphate Regulates Sorting Signal Recognition by the Clathrin-Associated Adaptor Complex AP2. Mol Cell. 2005;18: 519–531. doi:10.1016/j.molcel.2005.04.019

35. Mettlen M, Chen P-H, Srinivasan S, Danuser G, Schmid SL. Regulation of Clathrin-Mediated Endocytosis. Annu Rev Biochem. 2018;87: 871–896. doi:10.1146/annurev-biochem-062917-012644

36. Beck KA, Keen JH. Interaction of phosphoinositide cycle intermediates with the plasma membrane-associated clathrin assembly protein AP-2*. J Biol Chem. 1991;266: 4442–4447. doi:10.1016/S0021-9258(20)64342-3

37. Chandra M, Collins BM. The Phox Homology (PX) Domain. In: Atassi MZ, editor. Protein Reviews – Purinergic Receptors: Volume 20. Cham: Springer International Publishing; 2019. pp. 1–17. doi:10.1007/5584_2018_185

38. He K, Marsland III R, Upadhyayula S, Song E, Dang S, Capraro BR, et al. Dynamics of phosphoinositide conversion in clathrin-mediated endocytic traffic. Nature. 2017;552: 410–414. doi:10.1038/nature25146

39. Trey K. Sato, Michael Overduin, Scott D. Emr. Location, Location, Location: Membrane Targeting Directed by PX Domains | Science. 2001 [cited 20 May 2025]. Available: https://www.science.org/doi/10.1126/science.1065763?url_ver=Z39.88-2003&rfr_id=ori:rid:crossref.org&rfr_dat=cr_pub%20%200pubmed

40. Naslavsky N, Caplan S. The enigmatic endosome – sorting the ins and outs of endocytic trafficking. J Cell Sci. 2018;131: jcs216499. doi:10.1242/jcs.216499

41. Jovic M, Sharma M, Rahajeng J, Caplan S. The early endosome: a busy sorting station for proteins at the crossroads. Histol Histopathol. 2010;25: 99–112. doi:10.14670/hh-25.99

42. Singla A, Fedoseienko A, Giridharan SSP, Overlee BL, Lopez A, Jia D, et al. Endosomal PI(3)P regulation by the COMMD/CCDC22/CCDC93 (CCC) complex controls membrane protein recycling. Nat Commun. 2019;10: 4271. doi:10.1038/s41467-019-12221-6

43. Wallroth A, Haucke V. Phosphoinositide conversion in endocytosis and the endolysosomal system. J Biol Chem. 2018;293: 1526–1535. doi:10.1074/jbc.R117.000629

44. Christoforidis S, McBride HM, Burgoyne RD, Zerial M. The Rab5 effector EEA1 is a core component of endosome docking. Nature. 1999;397: 621–625. doi:10.1038/17618

45. Simonsen A, Lippe R, Christoforidis S, Gaullier J-M, Brech A, Callaghan J, et al. EEA1 links PI(3)K function to Rab5 regulation of endosome fusion. Nature. 1998;394: 494–498. doi:10.1038/28879

46. Odorizzi G, Babst M, Emr SD. Fab1p PtdIns(3)P 5-Kinase Function Essential for Protein Sorting in the Multivesicular Body - ScienceDirect. 1998 [cited 20 May 2025]. Available: https://www.sciencedirect.com/science/article/pii/S0092867400817079?via%3Dihub

47. Whitley P, Reaves BJ, Hashimoto M, Riley AM, Potter BVL, Holman GD. Identification of Mammalian Vps24p as an Effector of Phosphatidylinositol 3,5-Bisphosphate-dependent Endosome Compartmentalization*. J Biol Chem. 2003;278: 38786–38795. doi:10.1074/jbc.M306864200

48. Franco I, Gulluni F, Campa CC, Costa C, Margaria JP, Ciraolo E, et al. PI3K Class II α Controls Spatially Restricted Endosomal PtdIns3P and Rab11 Activation to Promote Primary Cilium Function. Dev Cell. 2014;28: 647–658. doi:10.1016/j.devcel.2014.01.022

49. Campa CC, Hirsch E. Rab11 and phosphoinositides: A synergy of signal transducers in the control of vesicular trafficking. Adv Biol Regul. 2017;63: 132–139. doi:10.1016/j.jbior.2016.09.002

50. Malek M, Kielkowska A, Chessa T, Anderson KE, Barneda D, Pir P, et al. PTEN Regulates PI(3,4)P2 Signaling Downstream of Class I PI3K. Mol Cell. 2017;68: 566–580.e10. doi:10.1016/j.molcel.2017.09.024

51. Román-Fernández Á, Roignot J, Sandilands E, Nacke M, Mansour MA, McGarry L, et al. The phospholipid PI(3,4)P2 is an apical identity determinant. Nat Commun. 2018;9: 5041. doi:10.1038/s41467-018-07464-8

52. Feng Z, Yu C. PI(3,4)P2-mediated membrane tubulation promotes integrin trafficking and invasive cell migration. Proc Natl Acad Sci U S A. 2021;118: e2017645118. doi:10.1073/pnas.2017645118

53. Balla T. Imaging and manipulating phosphoinositides in living cells. J Physiol. 2007;582: 927–937. doi:10.1113/jphysiol.2007.132795

54. Balla T, Várnai P. VISUALIZIATION OF CELLULAR PHOSPHOINOSITIDE POOLS WITH GFP-FUSED PROTEIN-DOMAINS. Curr Protoc Cell Biol Editor Board Juan Bonifacino Al. 2009;CHAPTER: Unit-24.4. doi:10.1002/0471143030.cb2404s42

55. Nasuhoglu C, Feng S, Mao J, Yamamoto M, Yin HL, Earnest S, et al. Nonradioactive Analysis of Phosphatidylinositides and Other Anionic Phospholipids by Anion-Exchange High-Performance Liquid Chromatography with Suppressed Conductivity Detection. Anal Biochem. 2002;301: 243–254. doi:10.1006/abio.2001.5489

56. Falasca M, Hamilton JR, Selvadurai M, Sundaram K, Adamska A, Thompson PE. Class II Phosphoinositide 3-Kinases as Novel Drug Targets. J Med Chem. 2017;60: 47–65. doi:10.1021/acs.jmedchem.6b00963

57. Kalasova I, Fáberová V, Kalendová A, Yildirim S, Uličná L, Venit T, et al. Tools for visualization of phosphoinositides in the cell nucleus. Histochem Cell Biol. 2016;145: 485–496. doi:10.1007/s00418-016-1409-8

58. Wills RC, Goulden BD, Hammond GRV. Genetically encoded lipid biosensors. Mol Biol Cell. 2018;29: 1526–1532. doi:10.1091/mbc.E17-12-0738

59. Hammond GRV, Ricci MMC, Weckerly CC, Wills RC. An update on genetically encoded lipid biosensors. Mol Biol Cell. 2022;33: tp2. doi:10.1091/mbc.E21-07-0363

60. Lemmon MA. Membrane recognition by phospholipid-binding domains. Nat Rev Mol Cell Biol. 2008;9: 99–111. doi:10.1038/nrm2328

61. Sato K, Norris A, Sato M, Grant BD. C. elegans as a model for membrane traffic. WormBook: The Online Review of C elegans Biology [Internet]. WormBook; 2018. Available: https://www.ncbi.nlm.nih.gov/books/NBK19650/

62. Lažetić V, Fay DS. Molting in C. elegans. Worm. 2017;6: e1330246. doi:10.1080/21624054.2017.1330246

63. Lažetić V, Fay DS. Conserved Ankyrin Repeat Proteins and Their NIMA Kinase Partners Regulate Extracellular Matrix Remodeling and Intracellular Trafficking in *Caenorhabditis elegans*. Genetics. 2017;205: 273–293. doi:10.1534/genetics.116.194464

64. Yochem J, Lažetić V, Bell L, Chen L, Fay D. *C. elegans* NIMA-related kinases NEKL-2 and NEKL-3 are required for the completion of molting. Dev Biol. 2015;398: 255–266. doi:10.1016/j.ydbio.2014.12.008

65. Joseph BB, Wang Y, Edeen P, Lažetić V, Grant BD, Fay DS. Control of clathrin-mediated endocytosis by NIMA family kinases. PLoS Genet. 2020;16: e1008633. doi:10.1371/journal.pgen.1008633

66. Joseph BB, Naslavsky N, Binti S, Conquest S, Robison L, Bai G, et al. Conserved NIMA kinases regulate multiple steps of endocytic trafficking. PLoS Genet. 2023;19: e1010741. doi:10.1371/journal.pgen.1010741

67. Lažetić V, Joseph BB, Bernazzani SM, Fay DS. Actin organization and endocytic trafficking are controlled by a network linking NIMA-related kinases to the CDC-42-SID-3/ACK1 pathway. PLoS Genet. 2018;14: e1007313. doi:10.1371/journal.pgen.1007313

68. Joseph BB, Blouin NA, Fay DS. Use of a Sibling Subtraction Method for Identifying Causal Mutations in Caenorhabditis elegans by Whole-Genome Sequencing. G3 GenesGenomesGenetics. 2017;8: 669–678. doi:10.1534/g3.117.300135

69. Milne SM, Edeen PT, Fay DS. TAT-1, a phosphatidylserine flippase, affects molting and regulates membrane trafficking in the epidermis of Caenorhabditis elegans. Genetics. 2024; iyae216. doi:10.1093/genetics/iyae216

70. Aung KT, Yoshioka K, Aki S, Ishimaru K, Takuwa N, Takuwa Y. The class II phosphoinositide 3-kinases PI3K-C2α and PI3K-C2β differentially regulate clathrin-dependent pinocytosis in human vascular endothelial cells. J Physiol Sci JPS. 2018;69: 263–280. doi:10.1007/s12576-018-0644-2

71. Domin J, Gaidarov I, Smith MEK, Keen JH, Waterfield MD. The Class II Phosphoinositide 3-Kinase PI3K-C2α Is Concentrated in the Trans-Golgi Network and Present in Clathrin-coated Vesicles*. J Biol Chem. 2000;275: 11943–11950. doi:10.1074/jbc.275.16.11943

72. Cheng S, Wu Y, Lu Q, Yan J, Zhang H, Wang X. Autophagy genes coordinate with the class II PI/PtdIns 3-kinase PIKI-1 to regulate apoptotic cell clearance in C. elegans. Autophagy. 2013;9: 2022–2032. doi:10.4161/auto.26323

73. Cheng S, Wang K, Zou W, Miao R, Huang Y, Wang H, et al. PtdIns(4,5)P2 and PtdIns3P coordinate to regulate phagosomal sealing for apoptotic cell clearance. J Cell Biol. 2015;210: 485–502. doi:10.1083/jcb.201501038

74. Liu J, Li M, Li L, Chen S, Wang X. Ubiquitination of the PI3-kinase VPS-34 promotes VPS-34 stability and phagosome maturation. J Cell Biol. 2018;217: 347–360. doi:10.1083/jcb.201705116

75. Zou W, Lu Q, Zhao D, Li W, Mapes J, Xie Y, et al. Caenorhabditis elegans myotubularin MTM-1 negatively regulates the engulfment of apoptotic cells. PLoS Genet. 2009;5: e1000679. doi:10.1371/journal.pgen.1000679

76. Lu N, Shen Q, Mahoney TR, Neukomm LJ, Wang Y, Zhou Z. Two PI 3-Kinases and One PI 3-Phosphatase Together Establish the Cyclic Waves of Phagosomal PtdIns(3)P Critical for the Degradation of Apoptotic Cells. PLOS Biol. 2012;10: e1001245. doi:10.1371/journal.pbio.1001245

77. Clancy JC, Vo AA, Myles KM, Levenson MT, Ragle JM, Ward JD. Experimental considerations for study of C. elegans lysosomal proteins. G3 GenesGenomesGenetics. 2023;13: jkad032. doi:10.1093/g3journal/jkad032

78. Frøkjær-Jensen C, Davis MW, Sarov M, Taylor J, Flibotte S, LaBella M, et al. Random and targeted transgene insertion in Caenorhabditis elegans using a modified Mos1 transposon. Nat Methods. 2014;11: 529–534. doi:10.1038/nmeth.2889

79. Yochem J, Tuck S, Greenwald I, Han M. A gp330/megalin-related protein is required in the major epidermis of Caenorhabditis elegans for completion of molting. Development. 1999;126: 597–606. doi:10.1242/dev.126.3.597

80. Shiwarski DJ, Darr M, Telmer CA, Bruchez MP, Puthenveedu MA. PI3K class II α regulates δ-opioid receptor export from the trans-Golgi network. Mol Biol Cell. 2017;28: 2202–2219. doi:10.1091/mbc.E17-01-0030

81. Ghosh RN, Mallet WG, Soe TT, McGraw TE, Maxfield FR. An Endocytosed TGN38 Chimeric Protein Is Delivered to the TGN after Trafficking through the Endocytic Recycling Compartment in CHO Cells. J Cell Biol. 1998;142: 923–936.

82. Roquemore EP, Banting G. Efficient Trafficking of TGN38 from the Endosome to the trans-Golgi Network Requires a Free Hydroxyl Group at Position 331 in the Cytosolic Domain. Mol Biol Cell. 1998;9: 2125–2144.

83. Paschinger K, Hackl M, Gutternigg M, Kretschmer-Lubich D, Stemmer U, Jantsch V, et al. A DELETION IN THE GOLGI α-MANNOSIDASE II GENE OF CAENORHABDITIS ELEGANS RESULTS IN UNEXPECTED NON-WILD TYPE N-GLYCAN STRUCTURES. J Biol Chem. 2006;281: 28265–28277. doi:10.1074/jbc.M602878200

84. Palmisano NJ, Meléndez A. Autophagy in *C. elegans* development. Dev Biol. 2019;447: 103–125. doi:10.1016/j.ydbio.2018.04.009

85. Leboutet R, Largeau C, Müller L, Prigent M, Quinet G, Rodriguez MS, et al. LGG-1/GABARAP lipidation is not required for autophagy and development in Caenorhabditis elegans. Zhang H, Kornmann B, editors. eLife. 2023;12: e85748. doi:10.7554/eLife.85748

86. Palmisano NJ, Meléndez A. Detection of Autophagy in Caenorhabditis elegans Using GFP::LGG-1 as an Autophagy Marker. Cold Spring Harb Protoc. 2016;2016: pdb.prot086496. doi:10.1101/pdb.prot086496

87. Philipp TM, Gong W, Köhnlein K, Ohse VA, Müller FI, Priebs J, et al. SEMO-1, a novel methanethiol oxidase in Caenorhabditis elegans, is a pro-aging factor conferring selective stress resistance. BioFactors. 2022;48: 699–706. doi:10.1002/biof.1836

88. Wang Y, Arnold ML, Smart AJ, Wang G, Androwski RJ, Morera A, et al. Large vesicle extrusions from C. elegans neurons are consumed and stimulated by glial-like phagocytosis activity of the neighboring cell. eLife. 2023;12: e82227. doi:10.7554/eLife.82227

89. Gillooly DJ, Morrow IC, Lindsay M, Gould R, Bryant NJ, Gaullier J-M, et al. Localization of phosphatidylinositol 3-phosphate in yeast and mammalian cells. EMBO J. 2000;19: 4577–4588. doi:10.1093/emboj/19.17.4577

90. Carvalho C, Moreira M, Barbosa DJ, Chan F-Y, Koehnen CB, Teixeira V, et al. ZYG-12/Hook’s dual role as a dynein adaptor for early endosomes and nuclei is regulated by alternative splicing of its cargo binding domain. Mol Biol Cell. 2025;36: ar19. doi:10.1091/mbc.E24-08-0364

91. Lemmon MA. Phosphoinositide Recognition Domains. Traffic. 2003;4: 201–213. doi:10.1034/j.1600-0854.2004.00071.x

92. Maekawa M, Fairn GD. Molecular probes to visualize the location, organization and dynamics of lipids. J Cell Sci. 2014;127: 4801–4812. doi:10.1242/jcs.150524

93. Lolicato F, Nickel W, Haucke V, Ebner M. Phosphoinositide switches in cell physiology - From molecular mechanisms to disease. J Biol Chem. 2024;300: 105757. doi:10.1016/j.jbc.2024.105757

94. Aggad D, Brouilly N, Omi S, Essmann CL, Dehapiot B, Savage-Dunn C, et al. Meisosomes, folded membrane microdomains between the apical extracellular matrix and epidermis. eLife. 2023;12: e75906. doi:10.7554/eLife.75906

95. Bae Y-K, Kim E, L’hernault SW, Barr MM. The CIL-1 PI 5-phosphatase localizes TRP Polycystins to cilia and activates sperm in C. elegans. Curr Biol CB. 2009;19: 1599–1607. doi:10.1016/j.cub.2009.08.045

96. Miao R, Li M, Zhang Q, Yang C, Wang X. An ECM-to-Nucleus Signaling Pathway Activates Lysosomes for C. elegans Larval Development. Dev Cell. 2020;52: 21–37.e5. doi:10.1016/j.devcel.2019.10.020

97. Li Y, Chen B, Zou W, Wang X, Wu Y, Zhao D, et al. The lysosomal membrane protein SCAV-3 maintains lysosome integrity and adult longevity | Journal of Cell Biology | Rockefeller University Press. 2016 [cited 20 May 2025]. Available: https://rupress.org/jcb/article/215/2/167/38758/The-lysosomal-membrane-protein-SCAV-3-maintains

98. Hermann GJ, Schroeder LK, Hieb CA, Kershner AM, Rabbitts BM, Fonarev P, et al. Genetic Analysis of Lysosomal Trafficking in Caenorhabditis elegans. Mol Biol Cell. 2005;16: 3273–3288. doi:10.1091/mbc.E05-01-0060

99. Miehe S, Bieberstein A, Arnould I, Ihdene O, Rütten H, Strübing C. The phospholipid-binding protein SESTD1 is a novel regulator of the transient receptor potential channels TRPC4 and TRPC5. J Biol Chem. 2010;285: 12426–12434. doi:10.1074/jbc.M109.068304

100. Jiang H, Sandoval Del Prado LE, Leung C, Wang D. Huntingtin-interacting protein family members have a conserved pro-viral function from Caenorhabditis elegans to humans. Proc Natl Acad Sci. 2020;117: 22462–22472. doi:10.1073/pnas.2006914117

101. Bannunah A, Cavanagh R, Shubber S, Vllasaliu D, Stolnik S. Difference in Endocytosis Pathways Used by Differentiated Versus Nondifferentiated Epithelial Caco-2 Cells to Internalize Nanosized Particles. Mol Pharm. 2024;21: 3603–3612. doi:10.1021/acs.molpharmaceut.4c00333

102. Joseph BB, Edeen PT, Meadows S, Binti S, Fay DS. An unexpected role for the conserved ADAM-family metalloprotease ADM-2 in Caenorhabditis elegans molting. PLoS Genet. 2022;18: e1010249. doi:10.1371/journal.pgen.1010249

103. Binti S, Melinda RV, Joseph BB, Edeen PT, Miller SD, Fay DS. A life cycle alteration can correct molting defects in Caenorhabditis elegans. Dev Biol. 2022;483: 143–156. doi:10.1016/j.ydbio.2022.01.001

104. Binti S, Linder AG, Edeen PT, Fay DS. A conserved protein tyrosine phosphatase, PTPN-22, functions in diverse developmental processes in C. elegans. PLoS Genet. 2024;20: e1011219. doi:10.1371/journal.pgen.1011219

105. Binti S, Edeen PT, Fay DS. Loss of the Na+/K+ cation pump CATP-1 suppresses nekl-associated molting defects. G3 GenesGenomesGenetics. 2024;14: jkae244. doi:10.1093/g3journal/jkae244

106. Stiernagle T. Maintenance of C. elegans. WormBook. 2006 [cited 23 Jan 2024]. doi:10.1895/wormbook.1.101.1

107. Montgomery MK. The Use of Double-Stranded RNA to Knock Down Specific Gene Activity. In: Miller WJ, Capy P, editors. Mobile Genetic Elements: Protocols and Genomic Applications. Totowa, NJ: Humana Press; 2004. pp. 129–144. doi:10.1385/1-59259-755-6:129

108. Farboud B, Meyer BJ. Dramatic Enhancement of Genome Editing by CRISPR/Cas9 Through Improved Guide RNA Design. Genetics. 2015;199: 959–971. doi:10.1534/genetics.115.175166

109. Farboud B, Severson AF, Meyer BJ. Strategies for Efficient Genome Editing Using CRISPR-Cas9. Genetics. 2019;211: 431–457. doi:10.1534/genetics.118.301775

110. Ghanta KS, Ishidate T, Mello CC. Microinjection for precision genome editing in *Caenorhabditis elegans*. STAR Protoc. 2021;2: 100748. doi:10.1016/j.xpro.2021.100748

111. Fay SF, Fay DS, Chhatre VE. CRISPRcruncher: A tool for engineering restriction sites into coding regions. MicroPublication Biol. 2021 [cited 23 Jan 2025]. doi:10.17912/micropub.biology.000343

112. Davis MW, Jorgensen EM. ApE, A Plasmid Editor: A Freely Available DNA Manipulation and Visualization Program. Front Bioinforma. 2022;2. doi:10.3389/fbinf.2022.818619

113. Redemann S, Schloissnig S, Ernst S, Pozniakowsky A, Ayloo S, Hyman AA, et al. Codon adaptation-based control of protein expression in C. elegans. Nat Methods. 2011;8: 250–252. doi:10.1038/nmeth.1565

114. Várnai P, Balla T. Visualization of phosphoinositides that bind pleckstrin homology domains: calcium- and agonist-induced dynamic changes and relationship to myo-[3H]inositol-labeled phosphoinositide pools. J Cell Biol. 1998;143: 501–510. doi:10.1083/jcb.143.2.501

115. Zhang L, Ward JD, Cheng Z, Dernburg AF. The auxin-inducible degradation (AID) system enables versatile conditional protein depletion in C. elegans. Dev Camb Engl. 2015;142: 4374–4384. doi:10.1242/dev.129635

116. Schindelin J, Arganda-Carreras I, Frise E, Kaynig V, Longair M, Pietzsch T, et al. Fiji: an open-source platform for biological-image analysis. Nat Methods. 2012;9: 676–682. doi:10.1038/nmeth.2019

117. Bolte S, Cordelières FP. A guided tour into subcellular colocalization analysis in light microscopy. J Microsc. 2006;224: 213–232. doi:10.1111/j.1365-2818.2006.01706.x

118. Dunn KW, Kamocka MM, McDonald JH. A practical guide to evaluating colocalization in biological microscopy. Am J Physiol - Cell Physiol. 2011;300: C723–C742. doi:10.1152/ajpcell.00462.2010

119. Fay DS. A biologist’s guide to statistical thinking and analysis. WormBook. 2013; 1–54. doi:10.1895/wormbook.1.159.1

